# *Syngap1* Promotes Cognitive Function through Regulation of Cortical Sensorimotor Dynamics

**DOI:** 10.1101/2023.09.27.559787

**Authors:** Thomas Vaissiere, Sheldon D. Michaelson, Thomas Creson, Jessie Goins, Daniel Fürth, Diana Balazsfi, Camilo Rojas, Randall Golovin, Konstantinos Meletis, Courtney A. Miller, Daniel O’Connor, Lorenzo Fontolan, Gavin Rumbaugh

## Abstract

Perception, a cognitive construct, emerges through sensorimotor integration (SMI). The genetic mechanisms that shape SMI required for perception are unknown. Here, we demonstrate in mice that expression of the autism/intellectual disability gene, *Syngap1*, in cortical excitatory neurons is required for formation of somatomotor networks that promote SMI-mediated perception. Cortical *Syngap1* expression was necessary and sufficient for setting tactile sensitivity, sustaining tactile object exploration, and promoting tactile learning. Mice with deficient *Syngap1* expression exhibited impaired neural dynamics induced by exploratory touches within a cortical-thalamic network known to promote attention and perception. Disrupted neuronal dynamics were associated with circuit-specific long-range synaptic connectivity abnormalities. Our data support a model where autonomous *Syngap1* expression in cortical excitatory neurons promotes cognitive abilities through assembly of circuits that integrate temporally-overlapping sensory and motor signals, a process that promotes perception and attention. These data provide systems-level insights into the robust association between *Syngap1* expression and cognitive ability.

## Introduction

Sensorimotor integration (SMI) refers to the neurophysiological phenomenon reflecting how sensory processing and motor output influence each other^1,2^. SMI is essential to a range of motor and higher cognitive functions, from posture, balance and movement control to attention, memory, and learning^3–7^. Sensory and motor signals are conveyed across multiple time scales through distributed networks and brain areas^5^. In rodents, disrupting SMI impairs neural representations of object features (texture, contour, and relative location), which are required for more complex cognitive functions to emerge, such as sensory perception and salience^8–12^. However, the neurobiological processes that shape the connectivity of distributed SMI networks that promote higher cognitive functions remain unknown. This hinders our understanding of the neural correlates of adaptive behaviors.

In addition to supporting a healthy brain, SMI processes are associated with a diverse range of disease/disorder states. This includes clumsiness, abnormal eye tracking, and altered sensory integration/reactivity, which are core features of neuropsychiatric disorders, such ASD and psychosis, and are also observed as “soft signs” in many neurological disorders ^13–18^. Genetic factors in the central and peripheral nervous systems have been implicated in abnormal sensory reactivity and altered motor control ^19–23^. Mutations in several genes have been identified in neurodevelopmental disorders (NDD) that feature alterations in sensory processing, motor control, and intellectual ability ^24–29^. Impaired SMI could, therefore, be a neural substrate of altered cognitive processes broadly observable in mental health disorders^28,30–33^. However, there have been comparatively few neurobiological investigations into how highly penetrant NDD risk genes contribute to SMI, resulting in a poor understanding of how gene expression shapes this essential neural process in health and disease.

We hypothesized that highly penetrant NDD genes regulate neurophysiological correlates of SMI required for higher cognitive functions. As an initial test of this hypothesis, we chose a relevant NDD gene and then tested how its expression contributed to SMI and associated cognitive functions. We chose *SYNGAP1*/*Syngap1* because expression of this NDD gene in humans and mice, respectively, is required for both sensory processing and motor control ^23,34^. Indeed, *de novo* mutations that lower *SYNGAP1* expression in humans cause a developmental and epileptic encephalopathy defined by impaired cortical excitability, postural/gait abnormalities, sensory processing impairments, and moderate-to-severe intellectual disability^35–40^. Moreover, a recent highly-powered genome-wide association study directly linked the *SYNGAP1* locus to cognitive abilities ^41^. Importantly, excellent mouse genetic tools are available for the study of *Syngap1* at the systems level. These tools enable region- and/or cell-specific bidirectional regulation of its expression, which allow spatial and temporal investigations into how *Syngap1* regulates distributed neural systems associated with SMI. These models have been used to uncover a role for *Syngap1* in cortical processing of sensory signals and control of motor responses required for decision-making ^23,40,42^. However, it remains unknown if *Syngap1* expression regulates neural correlates of SMI, and if so, how this contributes to constructs of cognition required for behavioral adaptation.

Here, *Syngap1* mouse genetic tools were used to explore how its expression regulates neurobehavioral correlates of SMI associated with constructs of cognition. To explore this, we utilized behavioral paradigms that rely on passive (receptive) and active (generative) whisker sensing to drive perceptual learning ^43,44^. In active tasks, tactile feedback enables closed loop, ongoing control of whisker motion, which promotes perception by enabling self-generated control of object exploration during tactile learning ^9,45^.The structural and functional connectivity of the rodent somatomotor whisker system has been extensively elucidated ^30,46–49^. The key nodes in higher-order whisker-related motor-sensory-motor (MSM) loops are known, and paradigms have been established that enable neurophysiological measurements of neuronal populations that mediate motor control during whisking, as well as tactile signals generated during object exploration. Importantly, disrupting self-generated motor control of whiskers during object exploration impairs perceptual learning ^9,50^. Thus, simultaneous tracking of whisker movement during object exploration and recording of activity within integrative neuronal populations enables elucidation of neurobiological principles that link SMI to cognition and behavior.

Using this framework, we utilized an array of *Syngap1* mouse models in learning paradigms that require the use of whiskers to generate percepts for behavioral adaptation. We paired these investigations with structural and functional analysis of somatomotor-associated neural circuits that integrate tactile and whisker motor signals. Combining these approaches, we demonstrate that *Syngap1* expression in cortical excitatory neurons is required for perceptual decision-making driven by tactile input, and for tactile-generated feedback control of whisker motion that underlies attention during active sensing. We also demonstrate that *Syngap1* regulates the structural/functional connectivity of cortical circuits within MSM loops known to integrate signals coding for touch and whisker motion. Together, these results demonstrate that a key function of *Syngap1* expression is to promote balanced integration of tactile and whisker motor signals within cortical sensorimotor loops. We propose that this form of abnormal SMI within the cortex of *Syngap1* mice contributes to reduced tactile sensitivity, poor attention, weak perceptual learning, and maladaptive behaviors. To our knowledge, this is the first demonstration for how impaired tactile processing associated with neurodevelopmental disorder genes can directly degrade cognitive performance required for behavioral adaptation.

## Results

### Syngap1 expression promotes whisker touch sensitivity and perceptual learning

We and others have previously demonstrated a role for *Syngap1* in both tactile learning and neural representations of tactile stimuli in somatosensory cortex ^23,51,52^. However, this past work did not define if, how, and to what extent *Syngap1* contributes to tactile learning through sensory processing. To begin to investigate sensory-mediated mechanisms linking *Syngap1* expression to perceptual learning underlying behavioral adaptation, we utilized a variation of a head-fixed tactile detection task where water-restricted animals were trained to provide a perceptual report of a passive whisker stimulation by licking a sensor that also supplies a water reward in mice older than 8 weeks of age. **(Fig. 1a)**. Three separate cohorts of *Syngap1^+/+^* (*wildtype* – normal SynGAP protein expression) and *Syngap1^+/−^* (+/−, germline heterozygous – half SynGAP protein expression) mice were trained, with each cohort receiving either a weak, medium, or strong whisker training stimulus during “Go” trials. Go trials were defined by a piezo deflection that induced a whisker stimulation; an animal scored a “hit” when licking the detector on these trials. “NoGo” trials were defined by a piezo deflection that did not translate into a whisker stimulus; an animal scored a “correct rejection” (CR) by withholding licking during these trials. We evaluated task performance of *wildtype* and *Syngap1^+/−^* mice by measuring total correct choices (hit on Go trials; CR on NoGo trials), overall Hit rate, overall False Alarm (FA) rate (FA = licking on a NoGo trial), and a trial discrimination index (*d’*). Stimulus intensity positively correlated with performance in *wildtype* mice, with faster learning over the 21-day training period with stronger stimulus intensity **(Fig. 1b)**, and improved trial discrimination at the end of training in strong versus weak training stimuli **(Supplementary Fig. 1a-i)**. However, *Syngap1* heterozygous mice exhibited deficient learning compared to controls as evidenced by fewer total correct choices, particularly in the strongest training stimulus **(Fig. 1b; Supplementary Table 1).** Additional analysis of trial data revealed that *Syngap1^+/−^* mice exhibited fewer Hits compared with littermate controls in the medium stimuli paradigm **(Supplementary Fig. 1d-f)**, and fewer hits with more FAs in the strong stimulus experiment **(Supplementary Fig. 1g-i)**. A generalized linear mixed model that considered all three cohorts revealed that the probability of correct choices in *Syngap1* heterozygous mice was less sensitive to increases in training stimulus intensity (F(57, 662.8)=2.8, p<0.001**, Supplementary Table 1)**. This suggested that *Syngap1* mice have reduced tactile sensitivity, though dual Hit and FA impairments, which could be a consequence of a fundamental disruption to distributed and generalized reinforcement learning mechanisms and/or motor control issues. To definitively determine if *Syngap1^+/−^* mice exhibit reduced tactile sensitivity, we carried out a “pull-back” experiment^53^ where animals that met acquisition criteria were subjected to a daily reduction in Go-stimulus intensity **(Fig. 1c)**. This experiment was possible because of a modified 3-step training paradigm that selected for a subset of *Syngap1*^+/−^ mice that learned to the same degree as *wildtype* littermates with the strong stimulus cohort **(Supplementary Fig. 1j-n)**. Thus, when additional *Syngap1^+/−^* mice were trained and the poor learners excluded from additional in-depth training, trial performance ended up no different between genotypes after 21 days of training **(Fig. 1d –** *first data point*). In this pull-back paradigm, there was an effect of genotype and an interaction between genotype and stimulus intensity in Go trials **(Fig. 1d)**. Indeed, in well-trained *Syngap1^+/−^* mice, Hit rates decreased faster relative to littermate controls as the stimulus intensity was reduced (F(4,40)=2.62, p<0.05). These head-fixed passive whisker stimulation task data demonstrate that *Syngap1* expression promotes tactile sensitivity.

**Fig. 1.**
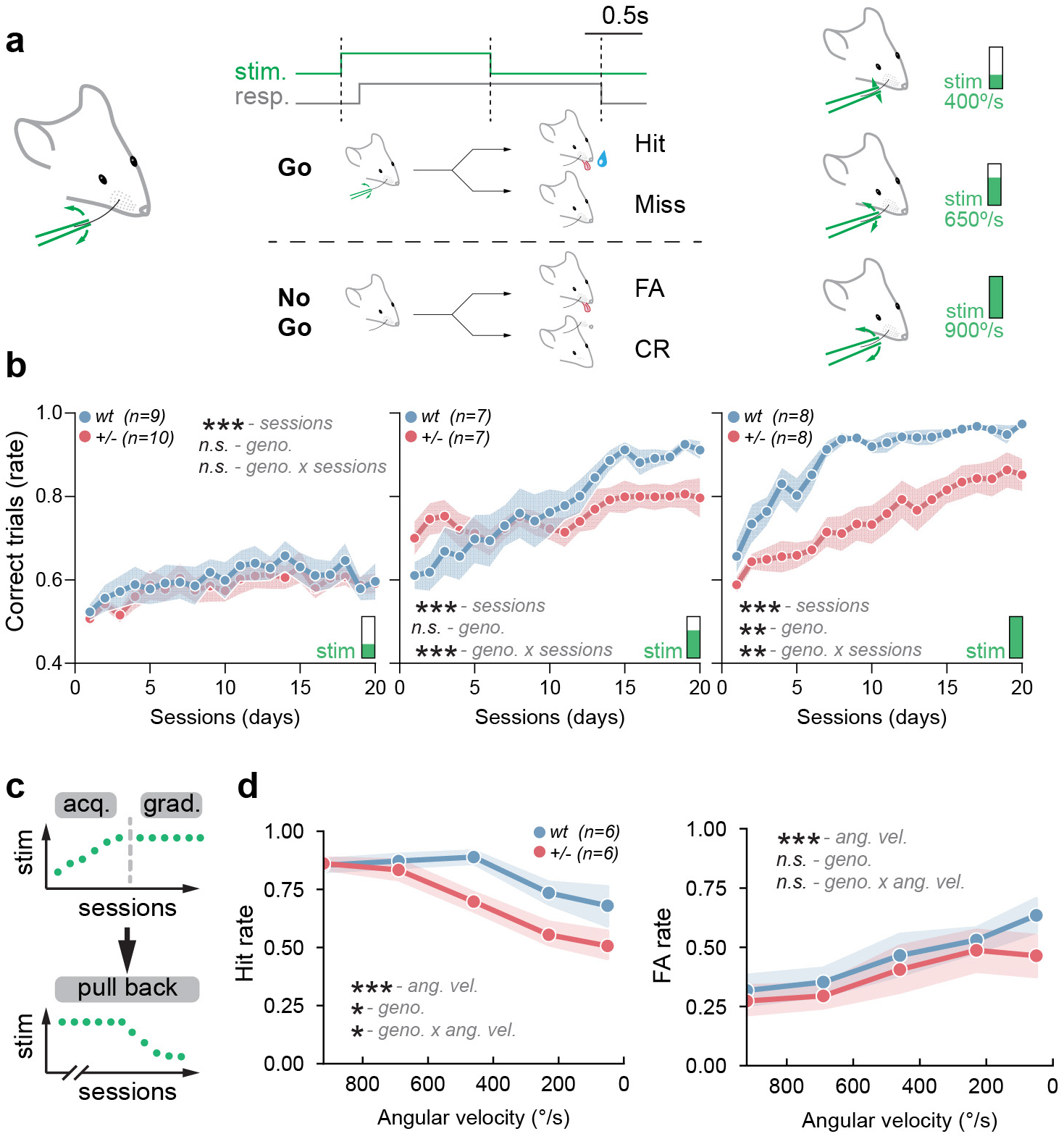
*Syngap1* expression promotes perceptual learning (tactile sensitivity) during passive tactile stimulus (WDIL). **a.** Schematic of the whisker-dependent instrumental learning paradigm (WDIL), including single whisker detection task structure and response outcomes with go trials being discriminated by single whisker deflection for 3 different whisker stimulus intensities (∼ 400, 650 and 900 °/s). During the WDIL task there was no auditory cue. The noise generated by the piezo during the Go and NoGo trial was masked by a 70dB white noise. The **b.** Fraction of total trials correct during WDIL for 3 different whisker stimulus intensities: ∼400°/s (wt: blue, n=9 and *Syngap1^+/−^*: +/−, red, n=10), 650°/s (wt: blue, n=7 and *Syngap1^+/−^*: +/−, red, n=7) and 900°/s (wt: blue, n=8 and *Syngap1^+/−^*: +/−, red, n=8). **c.** Summary schematic of the training phase and the reduced stimulation phase. **d.** False alarm (FA) and hit rates for animals that reached criteria and underwent the reduced stimulation phase (pull back) for wildtype (wt: blue, n=6) and *Syngap1^+/−^* (+/−: red, n=6). Number of animals per genotype is indicated in parentheses. Shaded areas represent the standard error of the mean (S.E.M.). p-value for main effects and interaction are indicated as n.s.: p>0.05, *: p<0.05, **: p<0.01, ***: p<0.001 detailed statistics are provided in Supplementary Table1.

This task reflects perceptual learning through a passive tactile stimulus. However, animals in the wild, including rodents and humans, most often acquire sensory information through self-generated movement of sense organs^54,55^. Therefore, we sought to determine the extent to which *Syngap1* expression regulated tactile sensitivity and associated perceptual adaptations in active sensing paradigms. First, we employed an active whisker-touch paradigm, **<u>N</u>**ovel **<u>O</u>**bjection **<u>R</u>**ecognition using only **<u>T</u>**exture **(NOR-T)**^23^, which was carried out in freely moving animals and was, therefore, ethological in nature. Freely moving mice in the dark were tasked with discriminating between two visually identical objects that only differed in texture **(Fig. 2a)**. Trimming whiskers in *wildtype* test mice prevented the expected shift in time spent around the novel textured object (whisker: t(9)=3.3, p=0.009; no whisker: t(9)=0.07, p=0.9; **Fig. 2b)**, confirming the task requires active use of whiskers. *Wildtype* mice could discriminate between the two objects, while *Syngap1^+/−^* mice could not (wildtype: t(7)=3.2, p=0.01; *Syngap1^+/−^*: t(5)=- 0.7, p=0.5; **Fig. 2c)**. However, when the difference in texture pattern density between the objects was greater (8 vs 5 instead of 9 vs 8 vertical ribs/cm, and presumably more perceptually salient, both genotypes could now discriminate (wildtype: t(9)=3.5, p=0.007; *Syngap1^+/−^*: t(9)=3.6, p=0.005; **Fig. 2d)**. Additional object recognition testing was conducted, which confirmed that poor texture discrimination in *Syngap1^+/−^* mice was caused by reduced tactile sensitivity rather than a more generalized impairment in brain function and behavior. For example, *Syngap1^+/−^* mice were able to discriminate equally well compared to littermate controls in a traditional novel object recognition task that robustly engages multisensory processes (wildtype: t(6)=2.9, p=0.02; *Syngap1^+/−^*: t(9)=4.3, p=0.002; **Fig. 2e)**. Together, these data demonstrate that *Syngap1* regulation of tactile sensitivity extends to texture discrimination.

**Fig. 2.**
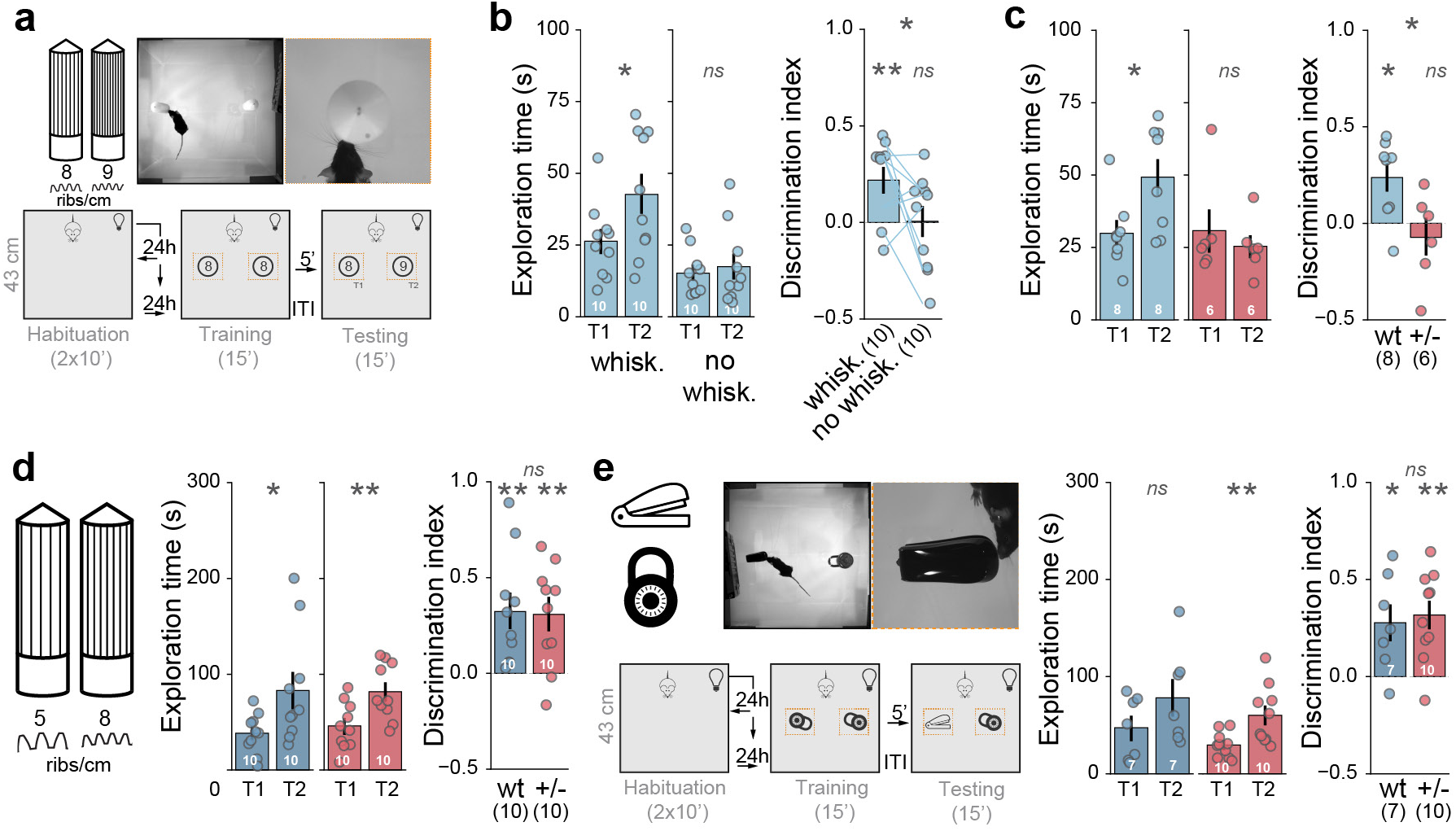
*Syngap1* expression promotes perceptual learning (tactile sensitivity) in ethologically relevant active sensing paradigms. **a.** Novel object recognition texture (NOR-T) task structure in which mice are tasked to discriminate between visually identical objects having either 8 or 9 verticals ribs/cm. The light level in the arena is 85 lux. **b.** Exploration time and discrimination index during the testing phase of the NOR-T for wild-type animals performing the task with (n=10) or without whiskers (n=10) for objects with 1 verticals ribs/cm difference. **c.** Exploration time and discrimination index during the testing phase of the NOR-T for wild-type (wt: blue, n=8) and *Syngap1^+/−^* (+/−: red, n=6) for objects with 1 verticals ribs/cm difference. **d.** Exploration time and discrimination index during the testing phase of the NOR-T for wild- type (wt: blue, n=10) and *Syngap1^+/−^* (+/−: red, n=10) for objects with 3 verticals ribs/cm difference. Mice with at least 10 sec of cumulative object exploration in the NOR-T were included in the analysis **e.** Novel Object Recognition (NOR) task structure with exploration time and discrimination index during the testing phase of the NOR for 2 different objects: a stapler and a lock in wild-type (wt: blue, n=7) and *Syngap1^+/−^* (+/−: red, n=10). Mice with at least 30 sec of cumulative object exploration in the NOR were included in the analysis. Number of animals per genotype is indicated in parentheses and in white within the bar graph. Error bars represent the standard error of the mean (S.E.M.). P-value are indicated as n.s.: p>0.05, *: p<0.05, **: p<0.01, ***: p<0.001. Detailed statistics are provided in Supplementary Table2.

In addition to texture discrimination, mice and rats actively use whiskers to perceive the location of objects relative to their head ^8,56,57^. To determine how *Syngap1* contributes to this form of tactile perception, we utilized a head-fixed Go/NoGo object localization task^58^. In this task, mice can use a single whisker to discriminate between two distinct object positions near the head **(Fig. 3a)**. Water-restricted animals were trained to discriminate between the Go and NoGo positions over ∼28 daily sessions. Correct choices on Go trials (e.g., licking the sensor) were reinforced with a water reward; FAs (licking on NoGo) resulted in a 15s timeout; misses (no licking on Go) went unrewarded and unpunished. While *Syngap1^+/−^* mice learned to detect the difference between the two locations, there was an effect of genotype, and an interaction between genotype and sessions, on the fraction of correct choices and in the trial discrimination index, *d’* **(Fig. 3b)**. To determine if impaired learning by mutants in this active tactile exploration task was related to impaired sensitivity to detecting object location, we again performed a pull-back experiment **(Fig. 3c)**. Statistical analysis revealed that performance was not the same between genotypes at the end of training (fraction of total correct: F(1,15) = 5.3, p=0.03). However, an interaction between genotype and object distance during pull-back sessions was detected in total correct choices (F(4, 60) = 2.6, p=0.04, **Fig. 3d)**. Consistent with impaired sensitivity (e.g., reduced precision) for detecting object location, the pull-back curve indicated that performance dropped off faster in *Syngap1^+/−^* mice compared to wildtype littermates.

**Fig. 3.**
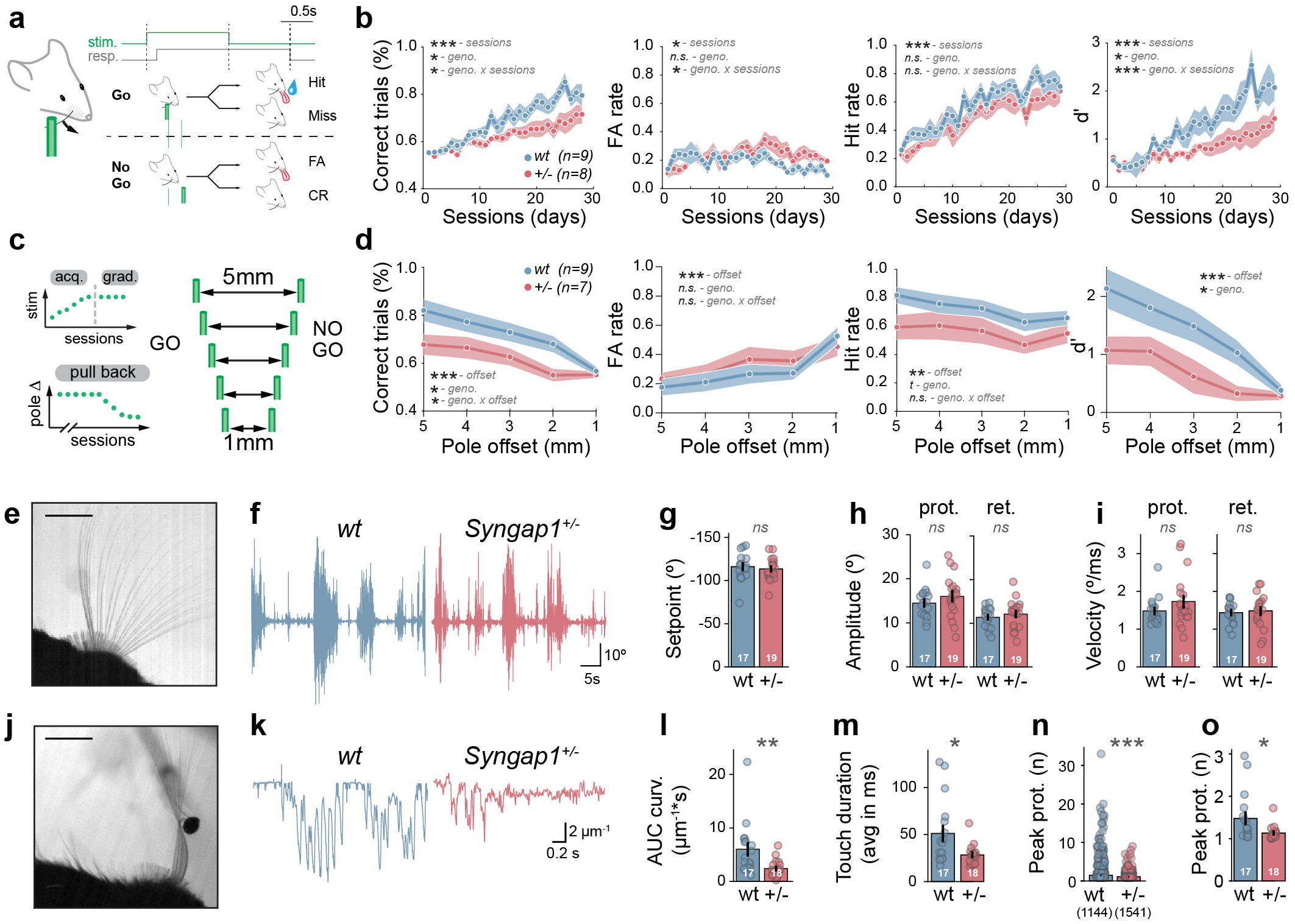
*Syngap1* expression promotes perceptual learning (tactile sensitivity) in a whisker- dependent discrimination task during instrumental learning based on pole location. **a.** Schematic of the behavioral paradigm during instrumental learning based on pole location. The pneumatic lift provided by the pneumatic linear slide also generates a sound that is not covered by the white noise and serves as an audio cue. **b.** Fraction of total trials correct, false alarm (FA) rate, Hit rate and d’ during the acquisition phase of the pole-location discrimination task. **c.** Schematic of the reduced stimulation phase of the pole-location discrimination task. Wildtype (wt: blue, n=9) and *Syngap1^+/−^* (+/−: red, n=8) **d.** Fraction of total trials correct, false alarm (FA) rate, Hit rate and d’ during the reduced stimulation phase of the pole-location discrimination task. Wildtype (wt: blue, n=9) and *Syngap1^+/−^* (+/−: red, n=7). Number of animals per genotype is indicated in parentheses. Shaded areas represent the standard error of the mean (S.E.M.). P-value for main effects and interaction are indicated as n.s.: p>0.05, *: p<0.05, **: p<0.01, ***: p<0.001. **e-i.** Whisker dynamics in free air and during object exploration. Superimposition of 40 frames representing whisking in wt mouse acquired at 500Hz illustrating free whisking of a single whisker, scale bar = 5mm **(e)**. Representative traces of whisker angle in wt (blue) and *Syngap1^+/−^* (red). Quantification of whisker setpoint **(g)**, amplitude **(h)** and velocity **(i)** of the protraction (prot.) and retraction (ret.) whisking phase during a 30 second recording window. **j-o.** Active touch dynamics of a single whisker for the first 30 seconds of pole presentation, superimposition of 40 frames representing active touch, scale bar = 5mm **(j)**. Representative traces of whisker curvature **(k)** and quantification of area under the curve (AUC) for whisker curvature **(l)**, average touch duration **(m)** and the total number of peak protractions detected for each individual touch event for wild-type (wt: blue, n=1144) and *Syngap1^+/−^* (+/−: red, n=1541) **(n)** and their respective animal average **(o)**. Error bars represent the standard error of the mean (S.E.M.) and number of animals for wild-type (wt: blue, n=17) and *Syngap1^+/−^*(+/−: red, n=19) is also indicated in white within the bar graph and/or in parentheses. P-value are indicated as n.s.: p>0.05, *: p<0.05, **: p<0.01, ***: p<0.001. Detailed statistics are provided in Supplementary Table3.

The motion of the whisker relative to an object is essential for determining object texture and location ^8,30,47,49^. Object contact causes whiskers to bend, eliciting torques and forces at the whisker base that are proportional to changes in whisker curvature. Strain within the follicle causes action potentials within trigeminal ganglion neurons. Therefore, reduced sensitivity for texture and location in *Syngap1* mice may be related to abnormal whisker motion during object exploration. To directly test this idea, we used a two-step approach that measured whisker kinematics in the same animals with and without the presence of a stationary pole **(Fig. 3e, j)**. Whisker dynamics were recorded using high-speed videography followed by offline location tracking with *WHISK*^58^ or DeepLabCut^59^ (**Supplementary Fig. 2**, **Video S1**). There was no effect of genotype on the setpoint (t(34)=0.5, p=0.6), maximum range of the whisker cycle (protraction: t(34)=1.1, p=0.3; retraction: t(34)=0.7, p=0.4), or whisker velocity during free air whisking (protraction: t(28.2)=1.4, p=0.2; retraction: t(34)=0.39, p=0.7, **Fig. 3f-i, Video S2)**, indicating that *Syngap1* expression does not regulate whisker kinematics in the absence of tactile input. We next quantified whisker dynamics in these same animals during whisking against a stationary pole **(Fig. 3j; Video S3)**, a paradigm that approximates the sensing process in the head-fixed pole localization task **(Fig. 3a)**. Physical interactions between the whisker and pole during rhythmic whisking induced whisker curvature during individual touch episodes **(Fig. 3j-k)**. Each episode of whisker contacting the object (i.e., touch episode) was extracted from the high-speed videos. Contact duration and whisker curvature for each touch episode was calculated for both genotypes. We observed that touch episodes generated smaller changes in whisker curvature in *Syngap1*^+/−^ mice compared to *wildtype* controls (AUC: U=57, p=0.001; **Fig. 3k-l)**. Moreover, there was an effect of genotype on touch duration, with *Syngap1^+/−^* mice exhibiting shorter touch durations than *wildtype* controls (t(18.8)=2.6, p=0.02; **Fig. 3m)**. Finally, we categorized touches based on how they influenced touch-induced pumps (TIPs), a specific type of whisker dynamic where the animal purposefully holds the whisker on the pole and engages in a “pumping” behavior once the pole is perceived ^60^. Importantly, TIPs elicit attention downstream of perception ^50^, and therefore this whisker kinematic provides a proxy measure of attention levels. We categorized all touches into four TIP categories based on amplitude and acceleration of the whisker while in contact with the pole **(Supplementary Fig. 3; Video S4**). In the category defined by >2 changes in amplitude and acceleration during pole contact, which includes the long-lasting touches with substantial levels of integrated curvature (and presumably long bouts of attention), we found that there were fewer of these touches in *Syngap1* heterozygotes compared to *wildtype* controls **(Fig. 3n-o, o**: U=239, p=0.01). Thus, this finding is consistent with impaired attention, reduced object exploration, and reduced tactile sensitivity in *Syngap1^+/−^* mice.

### Syngap1 expression within cortical excitatory neurons promotes perceptual learning, touch sensitivity, and touch-induced changes to whisker motion

Somatosensory systems are distributed throughout the brain and body. Thus, to gain mechanistic insight into the role of *Syngap1* expression on tactile sensing, we sought to identify the regional and cell-type origins of *Syngap1* expression sufficient to explain tactile phenotypes in this animal model. We hypothesized that *Syngap1* expression within higher-order brain areas may be sufficient to explain its role in both whisker dynamics and tactile learning. This theory was based on literature demonstrating that *Syngap1* is enriched in cortical areas ^61,62^, combined with separate literature indicating that touch engages top-down MSM loops, which dynamically tune whisker dynamics during sensing by signaling downward to brainstem motor neurons ^30,63–65^. To do this, we utilized two established *Syngap1* mouse lines that conditionally regulate the gene’s expression in cortical glutamatergic neurons (e.g., *EMX1*+ neurons). One line enables conditional heterozygosity within *EMX1+* neurons during the mid-embryonic period (*EMX1*-*Syngap1*-*OFF*), while the other embryonically re-activates *Syngap1* expression in a heterozygous null background selectively within the EMX1+ population ^66,67^ (*EMX1*-*Syngap1*-*ON*) **(Fig. 4a, Supplementary Fig. 4)**. We first assessed whisker kinematics **(Fig. 4b)**. No effect of genotype was observed in free air whisking measures in either model **(Fig. 4c-e, Supplementary Table4)**, which is consistent with a lack of phenotypes observed in germline (whole body) *Syngap1*^+/−^ mice **(Fig. 3h-l)**. However, during pole presentation **(Fig. 4f)**, *EMX1*-*Syngap1*-*OFF* heterozygous mice largely phenocopied altered touch-induced whisker kinematics originally observed in germline *Syngap1^+/−^* mice **(Fig. 4g-j**; **Fig. 3l-o)**. Touch episodes were shorter and generated less curvature compared to *wildtype* littermates (t(23.7)=3.3,p=0.003). In contrast, *EMX1*-*Syngap1*-*“ON”* heterozygous mice did not express touch-regulated whisker motion phenotypes found in the other two models **(Fig. 4g-j, g**:t(35.9)=- 0.4,p=0.7), even though *Syngap1* expression was only re-activated within EMX1+ glutamatergic cortical projection neurons (e.g. thalamic, cerebellar, brain stem areas, as well as the rest of the body, including all GABAergic neurons, remained heterozygous for *Syngap1* expression ^60,61^). This result demonstrates that *Syngap1* expression within EMX1+ neurons is necessary and sufficient for regulating touch-induced changes to whisker kinematics during pole exploration.

**Fig. 4.**
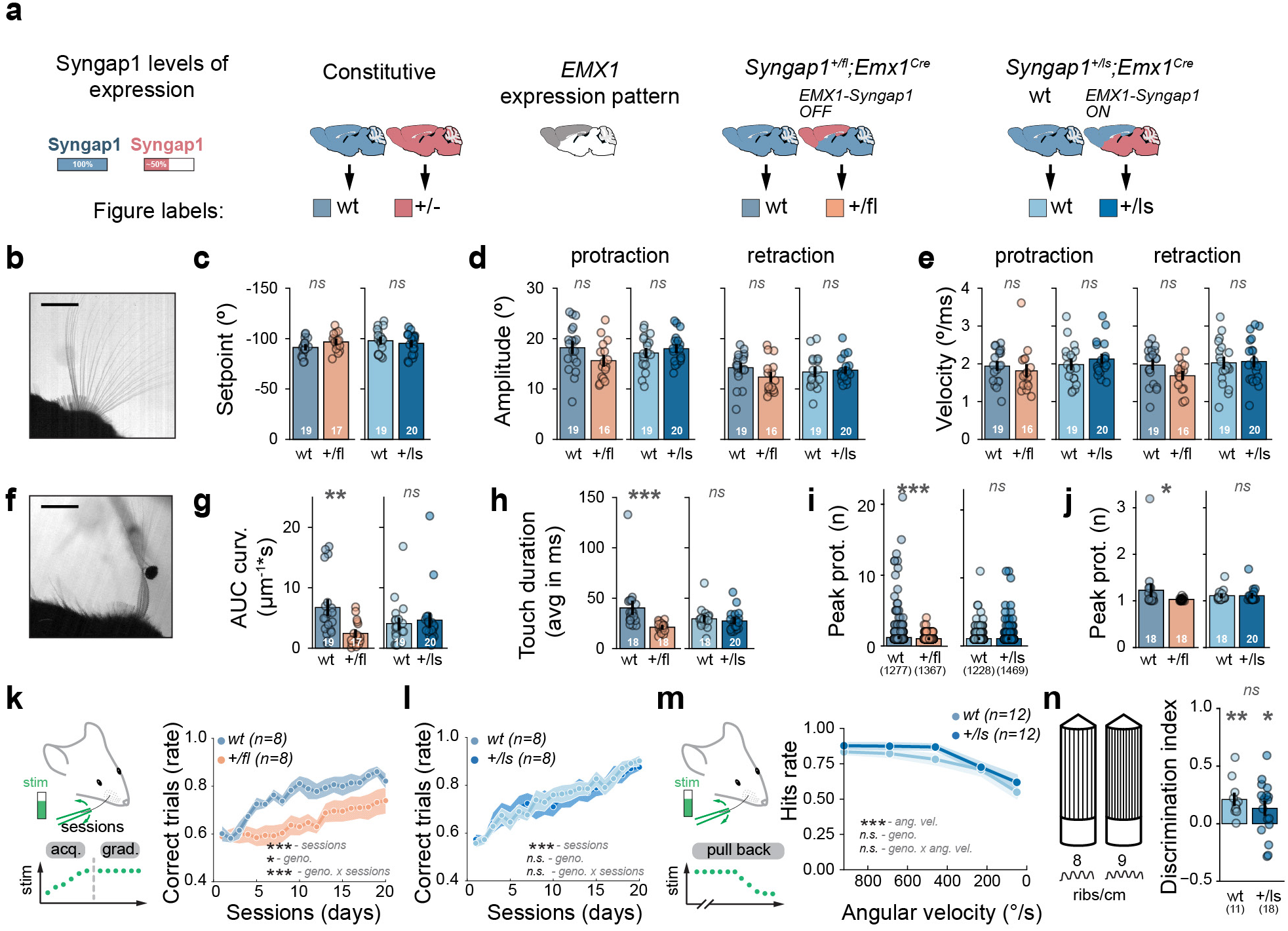
*Syngap1* expression within cortical excitatory neurons promotes tactile sensitivity and perceptual learning. **a.** Mouse line probing the impact of *Syngap1^+/−^* restricted to cortical excitatory neurons (*EMX1-Syngap1-OFF)*; cortical excitatory neuron rescue of *Syngap1^+/−^* (*EMX1-Syngap1-ON)* - all other cells in brain and body remain heterozygous for this gene. **b-e** Whisker dynamics in free air and during object exploration. Superimposition of 40 frames representing whisking in wt mouse acquired at 500Hz illustrating free whisking of a single whisker, scale bar = 5mm **(b)**. Quantification of whisker setpoint **(c)**, amplitude **(d)** and velocity **(e)** during the protraction (prot.) and retraction (ret.) whisking phase during a 30 second recording window for the *EMX1-Syngap1-OFF* mouse line (wt, n=19 and +/fl, n=17) *and EMX1-Syngap1-ON* mouse line (wt, n=19 and +/fl, n=20). **f-j** Active touch dynamics of a single whisker for the first 30 seconds of pole presentation, superimposition of 40 frames representing active touch, scale bar = 5mm **(f)**. Quantification of area under the curve (AUC) for whisker curvature **(g)**, average touch duration **(h)** and the total number of peak protraction detected for each individual touch event **(i)** and their respective animal average **(j)**. Color coding of groups is described in **(a)**. **k-n.** Modulation of *Syngap1* expression in cortical excitatory neuron population in tactile sensitivity assays. Summary schematic of the training phase during passive tactile stimulus (WDIL) performed with a whisker deflection of ∼650 °/s corresponding to a medium stimulus intensity and fraction of total trials correct in *EMX1-Syngap1-OFF* **(k)** and *EMX1-Syngap1-ON* **(l)** mouse lines. **m.** Hit rate for *EMX1-Syngap1-ON* animals that reached criteria and underwent the reduced stimulation phase of the WDIL performed with a whisker deflection of ∼650 °/s (pull back). **n.** Discrimination index during the testing phase of the NOR-T for objects with 10 groves spacing difference in *EMX1-Syngap1-ON* line. Mice with at least 10 sec of cumulative object exploration in the NOR-T were included in the analysis. Shaded areas and Error bars represent the standard error of the mean (S.E.M.) and number of animals for wild-type (wt: blue, n=17) and *Syngap1^+/−^*(+/−: red, n=19) is also indicated in white within the bar graph and/or in parentheses. P-value are indicated as n.s.: p>0.05, *: p<0.05, **: p<0.01, ***: p<0.001. Detailed statistics are provided in Supplementary Table4.

*Syngap1* expression within cortical glutamatergic neurons was also necessary and sufficient for promoting tactile sensitivity. For example, restricting *Syngap1* heterozygosity to cortical excitatory neurons phenocopied germline heterozygosity in the WDIL paradigm – there was a reduction in learning over the three-week training period in this task (**Fig. 4k, F**(19, 266) = 2.603, p<0.001**; Supplementary Fig. 5a**). In contrast, restricting *Syngap1* heterozygosity to all cells in the body except cortical glutamatergic neurons (i.e., EMX1-*Syngap1*-*ON*) resulted in no differences between genotypes in key measures of learning and trial discrimination (**Fig. 4l, F**(19, 133) = 0.5560, p=0.9**; Supplementary Fig. 5b).** Moreover, there was no effect of genotype in the pull-back portion of the study (**Fig. 4m, F** (5, 108) = 0.36, p=0.9**, Supplementary Fig. 5c),** demonstrating that detection sensitivity was normal in the EMX1+ rescue mice. Lack of phenotypes in the *Syngap1*-ON model was most likely driven by re-expression of *Syngap1* expression in the target neuron population (e.g., cortical excitatory neurons). This interpretation was supported by impaired pull-back sensitivity in non-Cre-expressing *Syngap1*-OFF mice compared to littermate controls **(Supplementary Fig. 5d-e)**. Indeed, these non-Cre expressing mice represent a distinct strain of *Syngap1* heterozygous knockout mice^66^. Thus, this result demonstrates reproducibility of the effect of *Syngap1* expression on tactile sensitivity. *EMX1-Syngap1-ON* mice were also tested in the NOR-T paradigm. Mice with *Syngap1* re-expressed in the EMX1+ population were able to discriminate between the two nearly identical textured objects (t(17)=2.4, p=0.03; **Fig. 4n, Supplementary Fig. 5f)**. Importantly, no unexpected germline deletion of LoxP sites^68^ was observed in the *EMX1-Syngap1-ON* mice **(Supplementary Fig. 4)**, demonstrating that regulation of *Syngap1* expression was indeed restricted to the expected target population. Together, these data demonstrate that expression of *Syngap1* within cortical glutamatergic neurons is both necessary and sufficient to produce touch-driven whisker control during object exploration, tactile sensitivity, and perceptual learning.

### Disrupted Neuronal Dynamics within the somatomotor network of Syngap1+/− mice

Given the autonomous role of *Syngap1* expression in cortex to sustain object exploration and associated perceptual learning, we hypothesized that neuronal dynamics within top-down MSM loops would be disrupted in *Syngap1^+/−^* mice. To directly test this, we measured unit spiking activity in higher-order whisker-related sensorimotor areas, including M1/M2, S1-BF, and whisker thalamus (VPM and POM areas), in *Syngap1* mice as they whisked in the presence or absence of a stationary pole. Neuronal spiking activity was recorded simultaneously across these regions in awake head-fixed *Syngap1^+/+^* and *Syngap1^+/−^* mice using multi-channel silicon probes **(Fig. 5a-d, Supplementary Fig. 6-8)**. During the two-hour recording period, lighting, auditory white noise, and the presence of a pole were varied to provide animals with a diverse sensory experience. After habituation, there was no difference in the amount of running on the treadmill during the recording **(Supplementary Fig. 8b)**. Across the three brain areas, there was no difference in the number of multiunit activity (MUA) clusters extracted from the two genotypes during electrophysiological recordings **(Supplementary Fig. 7b)**. In addition, there was no effect of genotype on the mean peak MUA spike rate in any of the three brain areas when activity was averaged for the entire recording period **(Fig. 5e)**. This suggests that the absolute level of ongoing spiking activity within the somatomotor network over prolonged time periods is not changed in head-fixed *Syngap1* mice.

**Fig. 5.**
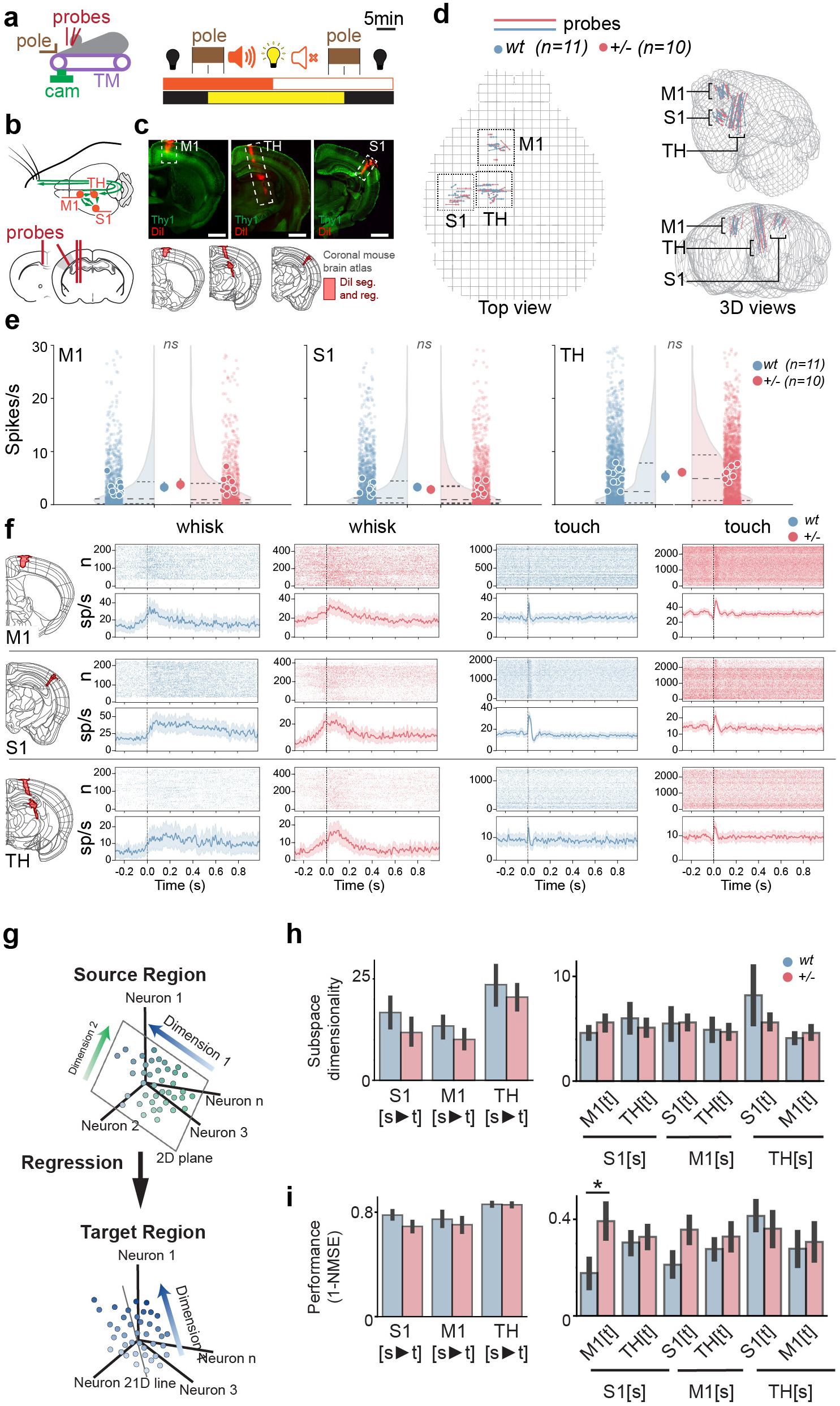
Impaired thalamocortical network dynamics during active sensing in *Syngap1^+/−^*mice. **a.** Experimental design and timeline during multi-channel silicon probe recordings. 3 probes were used during the recording along with camera and pole presentation while the mouse can move on a treadmill (TM). Different environmental variables were presented (light, sound and a pole that the mouse with a single whisker can interact with). **b.** Schematic of the neural circuitry and probe insertions during the experiment targeting key component of the somatosensory circuit (M1: motor cortex, S1: somatosensory cortex and TH: sensory thalamus). Post hoc analysis of electrode tracts did not have sufficient resolution to separate POM and VPM regions so the TH recordings may reflect units from both areas. **c.** Representative DiI (red) staining of probe implantation in motor cortex (M1), thalamus (TH), and somatosensory cortex (S1, left to right) with reconstructed DiI traces overlayed onto the Allen Brain atlas **d.** Multi probe track reconstruction in Allen Brain atlas for all wt (blue) and *Syngap1+/−* (+/−; red) for all the animals that underwent silicon probe recordings. **e.** Average spike rate (spikes/seconds) for extracted cluster including multiunit activity for both wt and *Syngap1+/−* (+/−) in M1, TH and S1. Small dots represent units, larger dots are animal means and largest dot represents group averages. Half violin plots illustrate the unit distributions. **f.** Raster and PSTH examples of firing for multi-units from M1 (top row), S1 (middle), TH (bottom) in wt and *Syngap1^+/−^* mice. **g-i.** Subspace analysis of neural communication in Syngap1+/− mice across and within cortical regions. **g**. Schematics of neural subspace dimensionality and predictive performance analysis across neural populations. High-dimensional neural activity in a source population is mapped onto a lower-dimensional subspace that is optimized for predicting the target population activity. **h**. The subspace dimensionality in the source population for predicting neural activity in the target population was determined by selecting the smallest number of dimensions at which the predictive performance, measured by 1-NMSE (Normalized Mean Squared Error), remained within one standard error of the mean (SEM) from the best performance. Bar plots display minimal subspace dimensionality with error bars representing SEM across different randomized choices of the source and target neural populations (see Methods). This analysis was applied across cortical regions, including within and between S1, M1, and TH. To evaluate the significance of differences in subspace dimensionality between genotypes a Monte Carlo permutation test was applied. A consistent trend of increased subspace dimensionality in Syngap1 could be observed, particularly in the connections from TH to S1 and S1 to M1, although these differences do not reach statistical significance. A similar trend was present during touches and inactive whisking epochs **(Supplementary Fig. 6)**. **c.** Predictive performance, measured as 1-NMSE (normalized mean squared error), of the target population activity based on the source population activity across the same-region and cross-region pairs.

Periods of free-air whisking and pole exploration represented only a fraction of the total time during the recording session. Therefore, we hypothesized that genotype-specific changes in neural activity would emerge during specified behavioral epochs defined by whisking with and without the pole present. Discrete free whisking (no pole) and touch (pole present) events were identified from high-speed video recordings **(Video S1).** These events represent two distinct behavioral transitions – from stationary to self-generated whisker movement **(whisking – VideoS2)** and from self-generated whisker motion to object contact inducing curvature **(touch – Video S3)**. We identified whisking and/or touch responsive MUA clusters in each brain area from both genotypes **(Fig. 5f)**. In animals from both genotypes, the dynamics of spike rate modulation were distinct in MUA clusters during free whisking compared to touch, and these patterns of activity agreed with past studies using similar recording techniques^69^. For example, free-air whisking units displayed prolonged activity on the order of hundreds of milliseconds, while touch-responsive units displayed activity for much shorter periods of time **(Video S5)**. Distinct unit modulation associated with free whisking and touch is consistent with the unique time scale of the two behaviors. Free whisking bouts were variable, though they usually lasted for hundreds of milliseconds and beyond^70^, while individual touches were generally an order of magnitude faster, with 80% of touches lasting less than 50ms **(Fig. 3m and 4h; Supplementary Fig. 8c)**.

The extraction of units modulated by whisker motion and/or touch allowed us to explore neural dynamics within and across the three regions of the somatomotor cortex as animals whisked with and without the pole present. We first quantified communication between neural populations by estimating the dimensionality of a subspace in the source population that best predicts activity in a target population^71^. We did this to test the hypothesis that *Syngap1* haploinsufficiency generally impairs the function of the somatomotor network during periods of active tactile exploration. Specifically, we identified a subspace within the source region’s neural activity that optimally predicts the activity in the target region. The dimensionality of this subspace serves as a measure of how many neural dimensions are involved in the communication between the two regions **(Fig. 5g)**. In *Syngap1^+/−^* mice, the analysis of neural communication across neural populations revealed a significant increase in predictive performance, particularly in cross-area interactions during whisking (**Fig. 5h-i**). A significant increase in 1-NMSE was observed specifically in the S1 to M1 region pair in mutants, indicating that the neural activity in S1 is more predictive of M1 activity compared to wildtypes. This enhanced predictive accuracy suggests that there may be altered functional connectivity between S1 and M1 in *Syngap1^+/−^* mice. The trend of increased predictive performance from S1 to M1 in *Syngap1* mutants during whisking was also consistent across behavioral epochs, though not all were statistically significant. This increased accuracy was not accompanied by statistically significant changes in subspace dimensionality, although there was a consistent trend toward greater dimensionality in the mutants, particularly in cross-regional interactions. This pattern was observed across different behavioral conditions, with a more pronounced effect during inactive epochs and a subtler, though present, trend during touch **(Supplementary Figure 6)**. These findings indicate that the *Syngap1* mutation may have a broad impact on the thalamocortical communication network, affecting both the strength and structure of inter-regional neural interactions.

### Features of neuronal spiking activity within nodes of the somatomotor network

To begin to understand how neuronal dynamics are altered in *Syngap1* mice, we next explored unit spiking properties across all three regions during whisking events with and without the presence of the stationary pole. Measuring spike rate modulation during the two distinct behaviors revealed numerous measures with genotype effects **(Fig. 6a, Supplementary Table 6)**. In general, motor signals associated with free whisking were significantly increased in *Syngap1^+/−^* compared to *Syngap1^+/+^* mice. Furthermore, the increase in motor activity during free whisking observed in M1 of *Syngap1^+/−^* mice was also present in S1 and thalamus, with thalamus resembling M1 **(Fig. 6a)**, while S1 MUAs demonstrated a less notable increase in activity compared to that observed in M1 and thalamus **(Fig. 6a)**. Together, these findings demonstrate that *Syngap1* mice have a generalized increase in whisker motor signals within MSM network nodes.

**Fig. 6.**
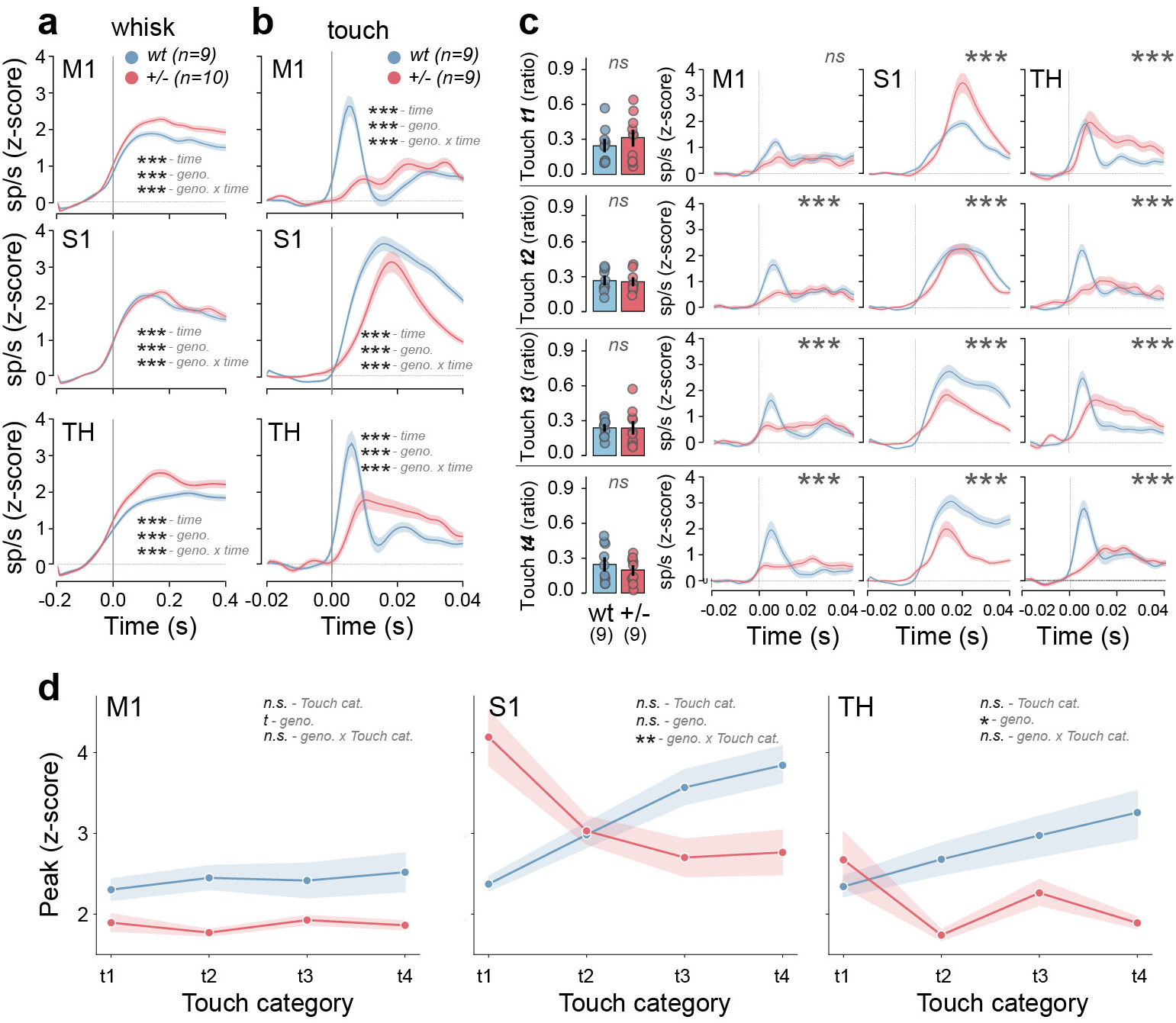
Impaired nodes of thalamocortical network during active sensing in *Syngap1^+/−^* mice. **a-b.** Averaged PSTHs for responsive clusters showing spike rates as z-scores during whisking onset **(a)** and touch **(b)** for M1, S1 and TH. **c.** Characterization of spike rates (as z-scores) from **(b)** based on strength of whisker curvature (t1: 0-25 percentile, t2: 25-50 percentile, t3: 50-75 percentile and t4: 75-100 percentile of maximum whisker curvature) for M1, S1 and TH in wt and *Syngap1^+/−^* mice and the corresponding peak amplitudes of z-score firing rates for each percentile of whisker touch **(d).** Shaded areas and Error bars represent the standard error of the mean (S.E.M.) and number of animals for wild-type (wt: blue, n=17) and *Syngap1^+/−^* (+/−: red, n=19) is also indicated in white within the bar graph and/or in parentheses. P-value are indicated as n.s.: p>0.05, *: p<0.05, **: p<0.01, ***: p<0.001.

In contrast, touch-related activity was generally decreased across these three regions in *Syngap1* mutants **(Fig. 6b)**. Analysis of MUA cluster dynamics during all touch events revealed significantly reduced peak spike rates in *Syngap1^+/−^* relative to *Syngap1^+/+^* mice. The general finding of reduced activity across the tactile-motor loop during touch in *Syngap1* mutants is consistent with our findings of reduced curvature of the whisker during individual object touches in animals with reduced expression of *Syngap1* **(Fig. 3l**; **Fig. 4g)**. In addition to abnormalities in peak spike rates, there was a clear disparity in the temporal onset of spiking in response to touch in mutants relative to wildtype littermate controls **(Fig. 6b)**. Unit dynamics in *Syngap1^+/−^* mice appeared to be low-pass filtered compared to wildtype littermates. In the thalamus, peak touch-dependent modulation was delayed, while duration of touch-related activity was prolonged in *Syngap1^+/−^* mice compared to wildtypes **(Fig. 6b)**. In M1/M2, a biphasic response was noted in wildtypes, with a rapid initial peak of activity occurring in response to object touch **(Fig. 6b)** and a secondary peak occurring 10-20ms later, which likely reflects recurrent activity arriving from other brain areas. In contrast, the initial peak after touch was reduced in *Syngap1^+/−^* mice, while the delayed responses appeared intact. Importantly, we observed reduced touch-induced spiking within S1-BF integrator units of *Syngap1* mutants, which responded to both whisker motion and pole-touch **(Supplementary Fig. 8e)**.

Reduced touch-related activity in units from *Syngap1^+/−^* mice could be a consequence of reduced psychomotor properties of the whisker during pole exploration (e.g., reduced average curvature during touch episodes). However, we have reported previously that controlled whisker curvature induced by passive whisker deflections results in reduced calcium-related neuronal activity in S1-BF from *Syngap1* mutants^23^. Thus, it is also possible that *Syngap1^+/−^* somatomotor networks may encode fewer spikes per unit of curvature. To resolve these two possibilities, we clustered all touch events into four categories based on integrated total whisker curvature in response to pole contact (e.g., “small”, “medium”, “large”, and “extra-large” levels of curvature). When categorized this way, there was no difference in curvature between the genotypes within any of the four touch categories **(Fig. 6c)**. As a result, touch-responsive units could now be compared between genotypes in the context of similar average levels of curvature. When unit activity was reanalyzed across the four touch categories in each genotype, surprising results were obtained. In wildtype mice, the expected positive correlation between curvature category and average peak spike rate in all three brain areas was present **(Fig. 6d)**. However, in *Syngap1^+/−^* mice, this relationship was essentially inverted. For example, there was a clear negative correlation between the magnitude of average curvature and average unit spike rate in both S1-BF and thalamus. Indeed, in S1-BF, touches that generated the greatest whisker curvature had the least amount of associated unit activity **(Fig. 6c-d)**. Taken together, touch elicits weak and temporally altered spiking activity in units from all three somatomotor areas in *Syngap1* mutants, while units that responded to whisking exhibited enhanced activity in these same areas.

### Circuit pathologies within higher-order somatomotor network hubs in *Syngap1* mice

Given the observed somatomotor network dysfunction in these mice, we hypothesized that dysfunction arises in part through abnormal structure and/or function of specific circuits connecting nodes within this network. We chose to probe inputs onto somatosensory cortex (S1) neurons for three reasons. First, abnormal connections through S1 were disproportionally identified in the communication subspace analysis **(Fig. 5g-i; Supplementary Figure 6)**. Second, neurons in this region are known to integrate motor and touch signals required for perception ^47,72,73^. Third, within S1, *Syngap1* regulates dendritic morphogenesis in deep neurons^74^ and developmental synaptic connectivity of feed-forward excitation in upper lamina neurons^23^, which could lead to circuit connectivity alterations capable of influencing neuronal dynamics within this cortical area. To test this idea, a cell-type specific rabies virus (RBV) monosynaptic retrograde labeling technique^75^ was performed to trace brain-wide synaptic connectivity onto L5 S1 neurons in *Syngap1^+/+^* and *Syngap1^+/−^* mice **(Fig. 7a-c)**. We achieved regional and neuronal subtype selectivity by crossing *Syngap1^+/−^* mice to RBP4-Cre mice and then injecting viral vectors directly into S1 of adult mice. Traced neurons within anatomically defined brain areas were registered and quantified^76^ **(Supplementary Fig. 9a-d)**. The location of helper and rabies virus injections within L5 S1 was similar across all animals of both genotypes and a similar number of sections from each group was analyzed **(Supplementary Fig. 9e-h)**. Importantly, no difference between genotypes was observed in the number of double labeled L5 S1 starter cells (t(12)=-0.6,p=0.6) and there was a similar number of inputs (t(12)=- 0.8,p=0.4); e.g., number of eGFP-only neurons outside L5 S1) relative to the starter cell population **(Supplementary Fig. 9i-j)**. This indicated that the total long-range synaptic connectivity onto L5 S1 neurons was not different between genotypes, which agrees with prior data demonstrating no alteration in spine density of L5 S1 at PND60 in *Syngap1^+/−^* mice^74^.

**Fig. 7.**
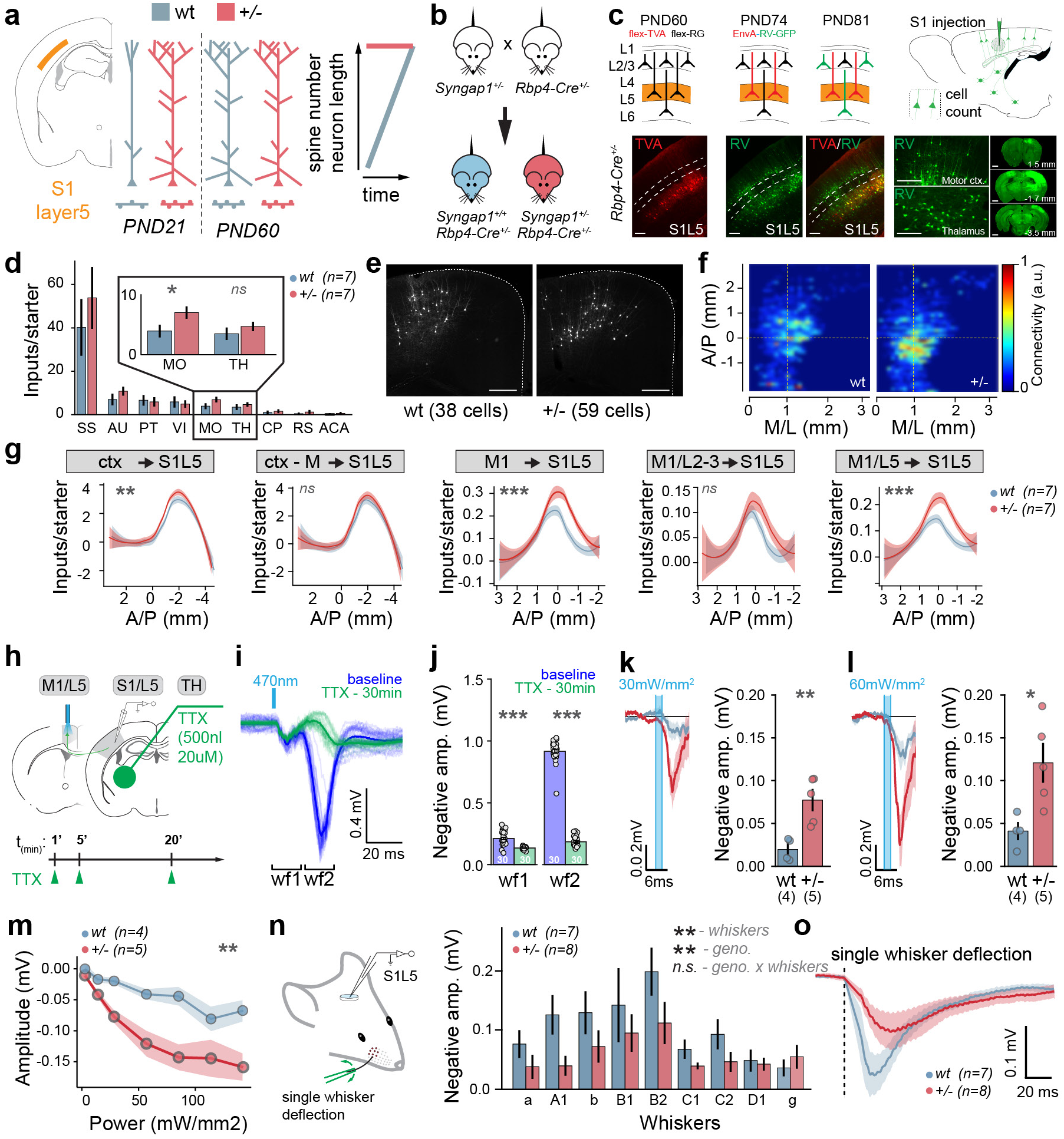
Specific components of the touch circuit within the somato-motor cortex are hyperconnected in *Syngap1^+/−^* mice. **a.** Model of accelerated maturation of S1L5 neurons based on prior published work^74^. **b.** Breeding strategy to enable S1L5 input mapping in wildtype and *Syngap1^+/−^*. **c.** Consecutive steps for cell type specific tracing: AAV transduction of Cre-dependent helper viruses (PND60, flex-TVA and flex-RG), rabies virus transduction (PND74, EnvA-RV-GFP), and rabies virus spread (PND81). Transduction of the helper virus flex-TVA and rabies virus are visualized respectively with the mCherry and GFP (red and green fluorescent signal) scale bar for brain section fluorescent images is 1mm and 100um for other images. On brain section the anterior-posterior (AP) coordinate relative to bregma in mm is indicated. **d.** Quantification of brain area specific inputs of S1L5 neurons normalized to the starter cell population (SS: somatosensory cortex, AU: auditory cortex, PT: posterior parietal association areas, VI: visual cortex, MO: motor cortex, TH: thalamus, CP: caudoputamen, RS: retrosplenial area, ACA: anterior cingulate area). **e.** Representative images of monosynaptic inputs from the motor cortex (+/+: wt, n=1 – 38 cells, +/−: *Syngap1^+/−^*, n=1 – 59 cells). Dotted line represents the brain outline on the motor cortex for coronal section at Bregma (AP=0mm). Scale bar = 200um. **f.** Flat map of motor cortex connectivity (arbitrary unit, a.u.) in antero-posterior (AP) and media-lateral (ML) averaged across all wt (n=7) and *Syngap1^+/−^*(+/−, n=7) mice. **g.** Quantification of the distribution of inputs to S1L5 normalized to the starter cell population in the antero-posterior axis for inputs from all cortical areas (ctx->S1L5), all cortical inputs after removing the motor inputs from the quantification (ctx – M -> S1L5), the motor cortex inputs (M1 -> S1L5), the L2/3 motor inputs (M1/L2-3 -> S1L5), L5 motor inputs (M1/L5 -> S1L5; blue: wt, n=7; red: *Syngap1^+/−^*:+/−, n=7; smoothed conditional means). **h.** Cartoon representing recording of LFPs during optogenetic stimulation of M1 an thalamic application of TTX resulting in the quantification of baseline and TTX LFP responses in S1L5. **e.** Field potential traces during baseline and 30 min post TTX bath application displaying 2 waveformes (wf1 and wf2). **j.** Quantification of the peak amplitude response during wf1 and wf2. **k-l.** Traces and peak LFP responses in S1L5 after M1L5 stimulation at 30 mW/mm2 **(k)** and 60mW/mm2 **(l)** stimulation intensities. **m.** Peak amplitude of LFP responses from different stimulation intensities in wt (n=5) and *Syngap1^+/−^* mice (n=4). **n.** Recording of LFPs during single whisker deflection. Individual peak LFP responses for each stimulated whisker. Nine whiskers (alpha, beta, A1, B1, B2, C1, C2 D1, gamma) were stimulated consecutively. **o.** LFP amplitude after single whisker deflection in wt (n=8) and *Syngap1^+/−^* (n=8) across 9 different whiskers. Shaded areas and Error bars represent the standard error of the mean (S.E.M.) and number of animals for wild-type (wt: blue, n=17) and *Syngap1^+/−^*(+/−: red, n=19) is also indicated in white within the bar graph and/or in parentheses. P-value are indicated as n.s.: p>0.05, *: p<0.05, **: p<0.01, ***: p<0.001. Detailed statistics are provided in Supplementary Table5.

We quantified relative connectivity onto L5 S1 neurons for all brain regions and observed a difference only for motor cortex (M1/M2) inputs (t(12)=-2.5, 0.03, **Fig. 7d-f)**. Grouping neurons within cortical origins across the antero-posterior axis revealed a genotype effect **(Fig. 7g**, Ctx- >S1/L5, F(1,13)=6.6,p=0.01). The genotype effect of afferent cortical connectivity onto L5 S1 neurons was driven by changes in motor cortex areas. Indeed, we observed an increase in eGFP-labeled neurons in M1/M2, but not other cortical areas, in *Syngap1^+/−^* mice compared to *wildtype* controls **(Fig. 7d-f)**. Moreover, when neurons originating from motor areas were removed from the “cortex” cluster, this difference was no longer significant **(Fig. 7g**, Ctx-M- >S1/L5, F(1,13)=6.6,p=0.01). Furthermore, the increased M1/2 labeling in *Syngap1^+/−^* mice was driven largely by neurons in deeper layers **(Fig. 7g**, M1/L2-3->S1/L5 and M1/L5->S1/L5). Thus, this unbiased screen of synaptic connectivity onto L5 S1 neurons revealed an increase in inputs arriving from motor cortex. Other connections may also be impacted in these mice, through these other regions did not reach levels of statistical significance.

To assess the validity of the significantly elevated motor-to-somatosensory cortex synaptic connectivity reported by RBV retrograde transsynaptic tracing, we measured the function of this input in *Syngap1* mice. An opto-probe was inserted into motor cortex of *Thy1*-ChR2 mice, which expresses ChR2 selectively within L5 neurons^77^. A single-channel electrode was lowered into L5 of S1 to record field potentials **(Fig. 7h).** Optogenetic activation within M1/M2 resulted in a biphasic waveform (WF1, WF2; **Fig. 7h-j)**. TTX injection into the thalamus had an outsized impact on WF2 compared to WF1 **(Fig. 7j)**. Given that WF1 occurred within a few milliseconds of the stimulus, these data together indicated that the early peak most likely reflects monosynaptic connections from M1/M2 to S1, while the later peak may reflect a multi-synaptic loop, such as the M1/2 > Thalamus > S1 loop. In a different set of experiments, a single-channel electrode was placed in L5 of S1 to record field potentials induced by ChR2+ in either wildtype or heterozygous *Syngap1* mice **(Fig. 7k-m)**. We observed an increase in WF1 in *Syngap1* mutants relative to littermate controls **(Fig. 7k-m**, at 60mW/mm^2^: t(5.6)=-3.4,p=0.01), which was consistent with the RBV retrograde tracing results. Moreover, long-range functional hyperconnectivity of M1/2 > S1 in *Syngap1* mice was not generalized, but was rather somewhat selective. In contrast to M1/M2 inputs to L5 of S1-BF, whisker deflections, which drive peripherally generated activity that arrives in S1 through thalamic feed-forward excitation, resulted in *<u>smaller</u>* synaptic responses in L5 S1 from *Syngap1^+/−^* mice compared to wildtype littermates (F(1,11)=16.2,p=0.002, **Fig. 7n-o)**. Together, these data demonstrate that L5 neurons in S1 of *Syngap1* mice receive a relatively strong input from M1/M2, a connection that relays whisker motor signals ^30,72^, but relatively weak afferent thalamocortical connectivity, an important connection that transmits whisker-touch signals into cortex. These circuit-specific impairments are consistent with abnormal dynamics within the somatomotor network as animals explore the environment using whiskers **(Fig. 5g-i).**

## Discussion

The molecular and cellular mechanisms that link SMI in higher brain regions to cognitive constructs that support adaptive behavior, such as attention and perception, are poorly understood. The motivation behind this study was to investigate the relationships between neurobiological correlates of SMI that promote cognitive functions and genetic factors associated with intellectual abilities. Here, we investigated how a gene strongly associated with sensory processing, cognitive ability, and autism risk, *Syngap1,* regulates sensorimotor processes in the cortex that support perception and attention. To understand how *Syngap1* contributes to SMI, we independently investigated how it contributes to sensory and motor processing underlying tactile sensing with whiskers. We also investigated the role of this gene directly in SMI through combined behavioral and electrophysiological observations during active whisker touch. Together, these data converged on a model where *Syngap1* promotes tactile perception and associated behavioral reactivity by assembling circuits that initially represent touch in the cortex. These circuits also integrate touch signals with whisker motor signals to promote an understanding of object features, such as location and texture, as well as eliciting attention. Our results indicate that *Syngap1* expression promotes assembly of cortical circuits that enable sensorimotor processing crucial for cognitive functions, including perception and attention. Thus, *SYNAGP1*/*Syngap1* haploinsufficiency may degrade intellectual ability by disrupting the function of a cortical sensorimotor network required to engage these cognitive processes.

Initially, we established that *Syngap1* loss-of-function leads to behavioral *<u>hypo-</u>*sensitivity when sensing with whiskers, which is expressed across tactile domains, including detection sensitivity, texture discrimination, and object localization **(Fig. 8a)**. These mouse studies are consistent with reports of tactile hyposensitivity and very high pain thresholds in *SYNGAP1-DEE* patients, including impaired behavioral reactivity in response to external stimuli applied to the body ^23,78^. Importantly, similar to the mouse model, this human patient population is defined by heterozygous loss-of-function variants within the *SYNGAP1* gene. Second, we found that *Syngap1* expression in cortical excitatory neurons is both necessary and sufficient for setting tactile detection thresholds that drive perception and associated adaptive behaviors **(Fig. 3)**. For example, well-trained *Syngap1* mutant mice made more perceptual errors relative to wildtype littermates in a subset of trials designed to measure tactile sensitivity. This suggested that poor learning in the tactile domain displayed by *Syngap1* mice may be explained, in part, through reduced whisker sensitivity. This phenotype was caused by its expression in cortical excitatory neurons because perceptual errors associated with whisker stimulus detection sensitivity were phenocopied in a mouse model with reduced *Syngap1* expression restricted to this population. Furthermore, when SynGAP protein was re-expressed in cortical excitatory neurons within *Syngap1* heterozygous mice, perceptual errors linked to whisker sensitivity were no longer present. A similar rescue of texture discrimination was observed in the *Syngap1* cortex-specific re-expression model. These findings, when integrated with past studies of other NDD risk factors, highlight the principle that individual genes can either promote or suppress neural mechanisms that dictate tactile sensitivity. Several NDD genes, such as *Shank3*, *Mecp2, and Fmr1* have been shown to regulate tactile sensitivity ^19,79^. However, for these risk factors, gene loss-of-function was linked to tactile *<u>hyper-</u>*sensitivity rather than *<u>hypo</u>*-sensitivity, which was explained by either cell-autonomous expression within the peripheral nervous system or in the CNS ^19,22,80–82^. Moreover, these studies focused on behavioral reactivity, and therefore did not link risk gene expression to neural correlates of cognitive constructs. The uniqueness of *Syngap1* with respect to tactile sensitivity in the context of these mouse studies is important because tactile hyposensitivity is also relatively common in NDD populations, including ASD ^26,80,83–86^. Thus, *Syngap1* mouse lines are emerging as genetic tools useful for elucidating the neural correlates linking NDD-associated sensory sensitivity to cognitive functions underlying adaptive behavior. In line with this, subsequent experiments in this study were aimed at understanding the potential neurobehavioral and neurophysiological correlates linking tactile hyposensitivity driven by *Syngap1* loss-of-function to impaired cognitive constructs.

**Fig. 8.**
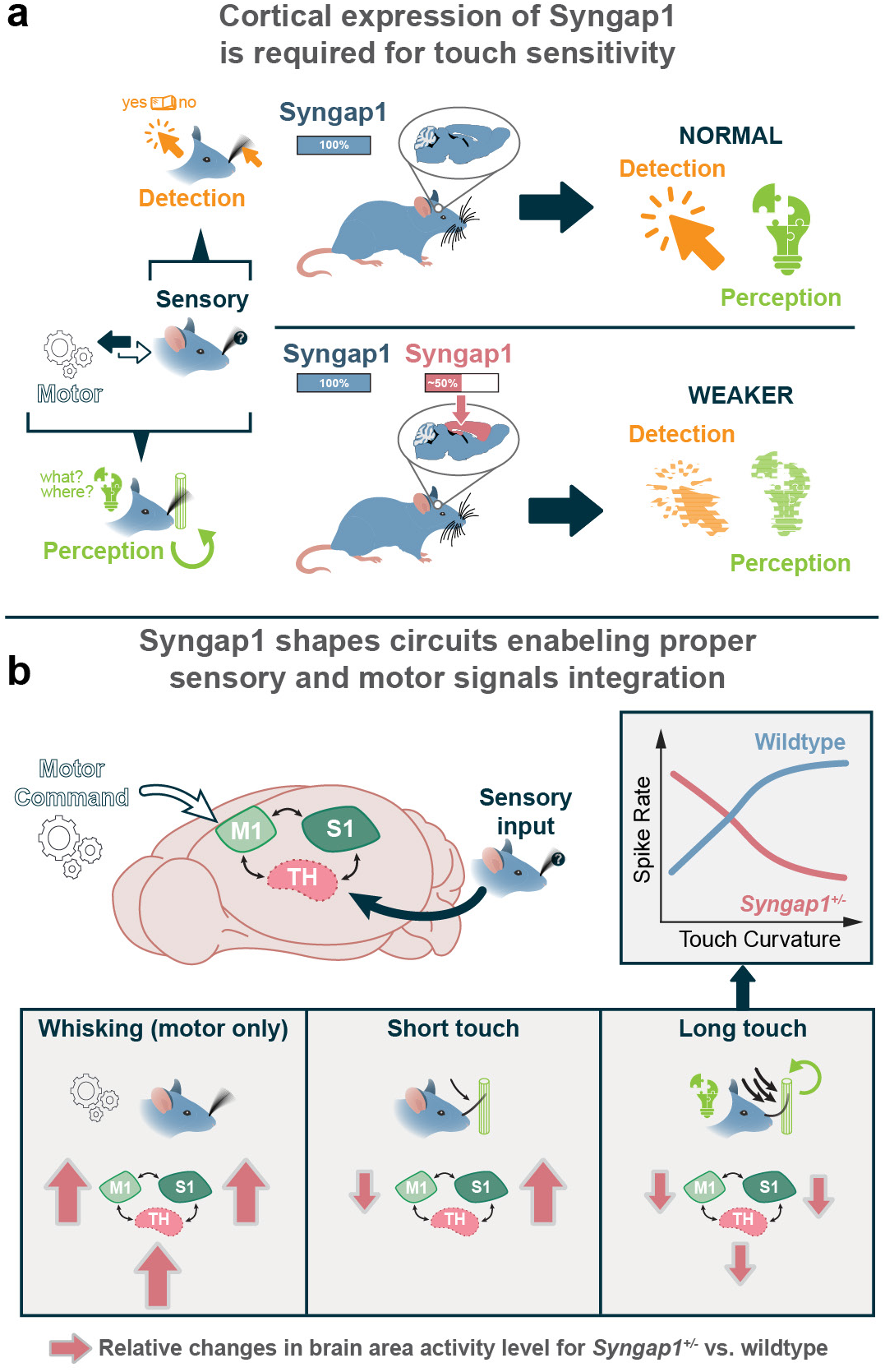
Model for how *Syngap1* expression shapes tactile sensitivity and perceptual behaviors through regulation of cortical SMI circuitry. **a.** When a *Syngap1+/−* animals explore objects with whiskers, touch information is poorly encoded in cortex, which prevents attentional processes from engaging a state-switch, which leads to reduced exploration times. Reduced exploration time will overtly decrease the opportunity for whiskers to code tactile signals required to compute object features. This impacts the strength of a tactile percept, which will impair perceptual learning and associated adaptive behaviors. **b.** In these mice, the touch signals that arrive in the cortex during the shortened object exploration are weakly encoded. In L5 somatosensory cortex, these weak touch signals are integrated with overly strong whisker motion (motor) signals, a SNR deficit that will degrade the ability of circuits to compute touch and texture features. Indeed, whisker motion and whisker touch generate two distinct streams of information at the level of the whisker follicle. The motor stream carries real-time whisker location information, while the touch stream contains a code for when touch occurred and how strong it was (through whisker curvature). These streams of information are integrated in barrel cortex L5 neurons, which enables an understanding of where the object rests relative to the head. This model can explain how *Syngap1* mice learn so poorly within the tactile domain. It is important to note that we also demonstrate that perceptual behaviors are disrupted when the task is biased toward whisker touch, but perception is intact for similar tasks when multisensory processes are available to the *Syngap1* animal.

Subsequent experiments led to a framework where *Syngap1* function within cortical glutamatergic neurons promotes tactile sensitivity through coding of touch signals in cortical somatomotor networks **(Fig. 8b)**. Several lines of evidence support this framework. *<u>First</u>*, impaired responses to passive whisker stimuli within cortical somatomotor areas of *Syngap1* mice is consistent with weak perceptual learning and reduced tactile sensitivity in the passive WDIL task. Our past studies have demonstrated that passive whisker deflections, the stimuli in WDIL, are weakly registered in barrel cortex of *Syngap1* mice, and this neurophysiological observation was dependent on *Syngap1* expression in the EMX1+ population ^23^. In that prior study, we noted that increased stimulus intensity of the whisker input in *Syngap1* mice caused larger deficits in barrel cortex activity compared to wildtype littermates. Whereas increasing stimulus intensity in wildtype mice led to larger cortical activation, this input/output relationship was impaired in *Syngap1* mice. Similarly, in this current study, we found that increasing the intensity of the stimulus during WDIL training induced more robust learning in wildtype mice. However, in *Syngap1* mutant mice, this was not the case. Increasing the stimulus intensity in these animals did not generate proportionally better learning as it did in wildtype, which led to qualitative differences in learning between genotypes **(Fig. 1)**. Thus, in these mice, there is a clear correlation between weak somatosensory cortex responses to passive whisker deflections and weak learning related to this same stimulus. Importantly, both behavioral reactivity to **(Fig. 4)**, and cortical representations of ^23^, the passive whisker stimulus are dependent upon *Syngap1* expression in cortical glutamatergic neurons.

*<u>Second,</u>* we observed abnormal touch-induced whisker exploratory behavior in *Syngap1* mice **(Fig. 2-3)**, including a reduction in the frequency and duration of touch-induced pumps (TIPs). This phenotype is also consistent with poor touch coding within cortical sensorimotor networks. Touch coding within higher-order motor-sensory-motor (MSM) loops is believed to generate touch-induced changes to whisker kinematics observed during active tactile sensing ^30,60,64,70^. Tactile signals representing initial object touch are believed to engage higher-order circuits that support constructs of cognition, such as attention ^60,70,87^. This leads to a shift in whisker sensing, with motion directed toward focused object exploration by the animal, which includes prolonged and purposeful object contacts that generate high levels of whisker curvature (e.g., TIPs). Thus, weak higher-order registration of initial object contacts in cortical networks is consistent with disrupted touch-induced whisker kinematics during object exploration. According to the existing model, touch signals in higher-order sensorimotor loops trigger prolonged object interactions by altering the function of whisker motor circuits in the brainstem, which is a neural correlate of attention ^30,64,65^. In support of this theory, we observed normal whisker kinematics in *Syngap1* mice during free-air whisking. Altered whisker kinematics were observed only when animals were presented with the stationary pole. This result would infer that the *Syngap1* expression in higher brain areas may be required to sustain object exploration by linking touch signals to attentional processes. This theory was additionally supported by the finding that *Syngap1* expression within the cortical excitatory neuron population is both necessary and sufficient for touch-induced changes to whisker kinematics **(Fig. 4)**. Therefore, *Syngap1* expression likely regulates neurobiological processes within this restricted neuronal population to promote integration of touch signals into attentional circuits that modulate whisker motor neurons in the brainstem. One possible cellular mechanism may involve the regulation of connectivity and/or function of cortical excitatory neurons that both code for touch and signal to brainstem neurons that drive whisker motion. This possibility was supported through our electrophysiological observations of reduced touch responses in units within higher-order somatomotor loops from *Syngap1* mutants **(Fig. 5)**. Future studies will be necessary to determine the extent to which *Syngap1* expression regulates touch-responsive neurons in the cortex that project to brainstem whisker motor circuits.

*<u>Third</u>*, we observed weak touch responses during pole exploration across higher-order somatomotor areas in *Syngap1* mice, including somatosensory and motor cortex, as well as whisker areas in thalamus **(Figs. 5-6)**. In these mutant animals, weak touch responses were present in the context of an increase in temporally-overlapping whisker motor signals. This sensorimotor imbalance reflected a real change in the signal-to-noise ratio of neurons that integrate touch with whisker motion. Importantly, we observed weak whisker touch responses in neurons from the barrel region of somatosensory cortex that also respond to whisker motion. This is a direct electrophysiological demonstration of altered SMI within the cortex of *Syngap1* animals. Reduced signal-to-noise for touch/motor integration is consistent with impaired object localization. Object localization is thought to be computed, in part, through convergent whisker touch and motor signals that are integrated within cortical sensory and motor areas ^30^. Electrophysiological correlates of reduced touch coding, as well as correlates of enhanced motor signals, were supported by circuit tracing and subsequent functional circuit validation studies in *Syngap1* mice. *Syngap1* mice had hyper-functional inputs from motor cortex that project to Layer 5 somatosensory cortex **(Fig. 7)**. These mice also possess a weak subcortical input that transmits touch signals into L5 somatosensory cortex.

It is unclear how reduced *Syngap1* expression in mice can cause impaired cortical touch representations, while also causing enhanced whisker motor representations. Understanding this dichotomy will provide insight into the neural correlates of impaired sensorimotor processing in NDDs. *Syngap1* expression may exert unique cell-autonomous functions within neuronal subtypes that comprise the EMX1+ population. Unique, cell-autonomous functional control over expression of neuronal features that directly assemble and refine circuits could lead to unpredictable circuit-specific impairments, not unlike what was observed in this current study. This hypothesis is supported by evidence in past studies demonstrating that *Syngap1* potently regulates cellular substrates of circuit assembly and refinement, such as dendritic morphogenesis, synaptic maturation, and cell/circuit-level forms of neural plasticity ^88^. Intriguingly, *Syngap1* regulates circuit-building substrates in a cell- and region-specific manner, sometimes in opposite directions. For example, it can both promote^74^ and constrain^23^ dendritic morphogenesis within distinct EMX1+ neuronal subtypes during developmental critical periods. Pertinent to this current study, the developmental maturation of L5b tufted neurons in somatosensory cortex is greatly accelerated in *Syngap1* mutant mice. These *Syngap1*-deficient neurons reach adult-like features of maturation weeks before similar neurons in wildtype littermates ^74^. This accelerated maturation spans major morphogenic developmental milestones, including accelerated dendritic differentiation, early acquisition of dendritic spines, and subsequent precocious spine pruning. In contrast, in this same model, and within the same brain area (somatosensory cortex), the development of L2/3/4 upper-lamina neurons feature arrested development ^23^. Altered maturation of neurons in cortex is not restricted to somatosensory cortex ^74^. De-synchronization of postmitotic dendritic and synaptic maturation across distinct neuronal subtypes and regions is consistent with impaired somatosensory network function observed in this study.

Significant mechanistic questions remain unanswered. One such area of exploration is to understand to what extent somatomotor system organization, such as local excitatory and inhibitory microcircuit connectivity or long-range connectivity among network hubs, is dictated directly (e.g., cell-autonomously) by *Syngap1* expression within distinct cellular subtypes. The challenge will be to then contrast this with indirect (e.g., non-cell autonomous/homeostatic) compensatory consequences that are engaged because of the direct cell-autonomous changes to neuronal structure and function downstream of *Syngap1* heterozygosity. This framework may help to explain the unexpected current result that M1/M2 excitatory projections to S1 exhibited outsized levels of hyperconnectivity compared to excitatory inputs originating from other brain areas. With regards to the cellular building blocks of cortical circuits, they can be broken down into nested categories, including neuronal vs. non-neuronal cells, excitatory vs. inhibitory neurons, or subclasses of excitatory (PT, IT) and inhibitory (PV, SOM) neurons. The current study provides evidence that cell-autonomous expression of *Syngap1* within cortical excitatory neurons is both *<u>necessary</u>* and *<u>sufficient</u>* for touch perception and touch-associated behavioral adaptations that require perception and attention. How these behavioral phenotypes relate to circuit assembly errors among different subclasses of excitatory neurons remains unknown. For example, accelerated maturation of L5 somatosensory cortex neurons^74^ in *Syngap1^+/−^* mice could be a non-cell-autonomous compensatory process driven by arrested development and hypofunction of upper lamina excitatory neurons in the same cortical area^23^.

It is also possible that cell-autonomous functions of *Syngap1* within inhibitory neurons may also play a role in the organization of the somatomotor system. Kwon et al^52^ recently reported that conditional knockdown of *Syngap1* within inhibitory neurons caused an impairment in performance during a similar whisker-dependent Go/NoGo task. However, genotype effects were detected for False Alarm responses, but not Hit responses. Because False Alarms may reflect impulsive responding, while Hits reflect a learned association between the whisker stimulus and the reward, these past results suggest that expression of *Syngap1* within inhibitory neurons may be restricted to motor control processes. Moreover, that study did not assess sufficiency of *Syngap1* within the inhibitory population. In our current study, manipulating SynGAP protein expression in cortical excitatory neurons essentially phenocopied observations in the germline heterozygous model. Integrating results from this past study with our current results indicate that there are autonomous roles for *Syngap1* in both excitatory and inhibitory cortical neurons, though whisker touch perception and attention is largely shaped by the function of the gene within the excitatory population. Regarding non-neuronal cell types, SynGAP is expressed in rodent astrocytes, but to lesser extent compared to neurons^51^. There are reports of sparse Cre activity in isolated astrocytes in the *Emx1*-Cre mouse line^67^, so it is theoretically possible that *Syngap1* expression in astrocytes could contribute to phenotypes observed in this study. However, we have empirically observed only a handful of glia that express a Cre reporter cassette in the EMX1-Cre mouse line (G.R., *unpublished observations)*, suggesting that cell-autonomous functions of *Syngap1* in astroglia may not substantially contribute to somatomotor/perceptual phenotypes. Future studies leveraging intersectional genetic perturbations that target *Syngap1* expression within defined cellular subtypes, across various brain areas, and across developmental epochs, will be necessary to tease out the direct vs. indirect effects of *Syngap1* on cellular elements that build top-down somatomotor network loops. Furthermore, given the recent genome-wide linkage of *SYNGAP1* to cognitive function^41^, it will be important determine to what extent *Syngap1* gain-of-function in animal models^89^ can enhance cognitive functions, and if so, how this might be achieved through altering neuronal dynamics within sensorimotor networks.

## Data availability

The data that support the findings of this study are available from the corresponding author upon reasonable request.

## Code availability

Custom code was written in Matlab, R and Python for data analysis. Codes which are not already available in public repository (see Methdods) are available upon reasonable request.

## Acknowledgements

This study was supported by funding provided by NIMH (MH096847) and NINDS (NS110307) to G.R./C.A.M and G.R. (w/D.O. as CO-I), respectively. We thank Gogce C. Crynen, PhD, Director of the UF Scripps Biostatistics Core, for consultation on appropriate application of statistical models to complex data sets. L. F. is supported by funding from Excellence Initiative of Aix Marseille Universit^é^ - A*MIDEX (Turing Centre for Living Systems). We would like to thank Dakota Roberts, Stephanie A. Miller and Izan Gonzalez Fayos for technical assistance with histology and tissue processing.

## Author Contributions

T.V and S.D.M. performed experiments, designed experiments, analyzed data, co-wrote the manuscript, and edited the manuscript. T.C. performed experiments, designed experiments, analyzed data, and edited the manuscript. D.F. and D.B. analyzed and interpreted data. L.F. designed experiments, analyzed data, interpreted data, and edited the manuscript. C.R. performed experiments, designed experiments, and generated reagents. R.G. designed experiments and generated data. K.M. provided key reagents and interpreted data. C.A.M secured funding, designed experiments, interpreted data, and edited the manuscript. D.O. designed experiments, interpreted data, edited the manuscript. G.R. conceived the study, secured funding, designed experiments, interpreted data, co-wrote the manuscript, and edited the manuscript.

---

**Supplementary Fig. 1.**
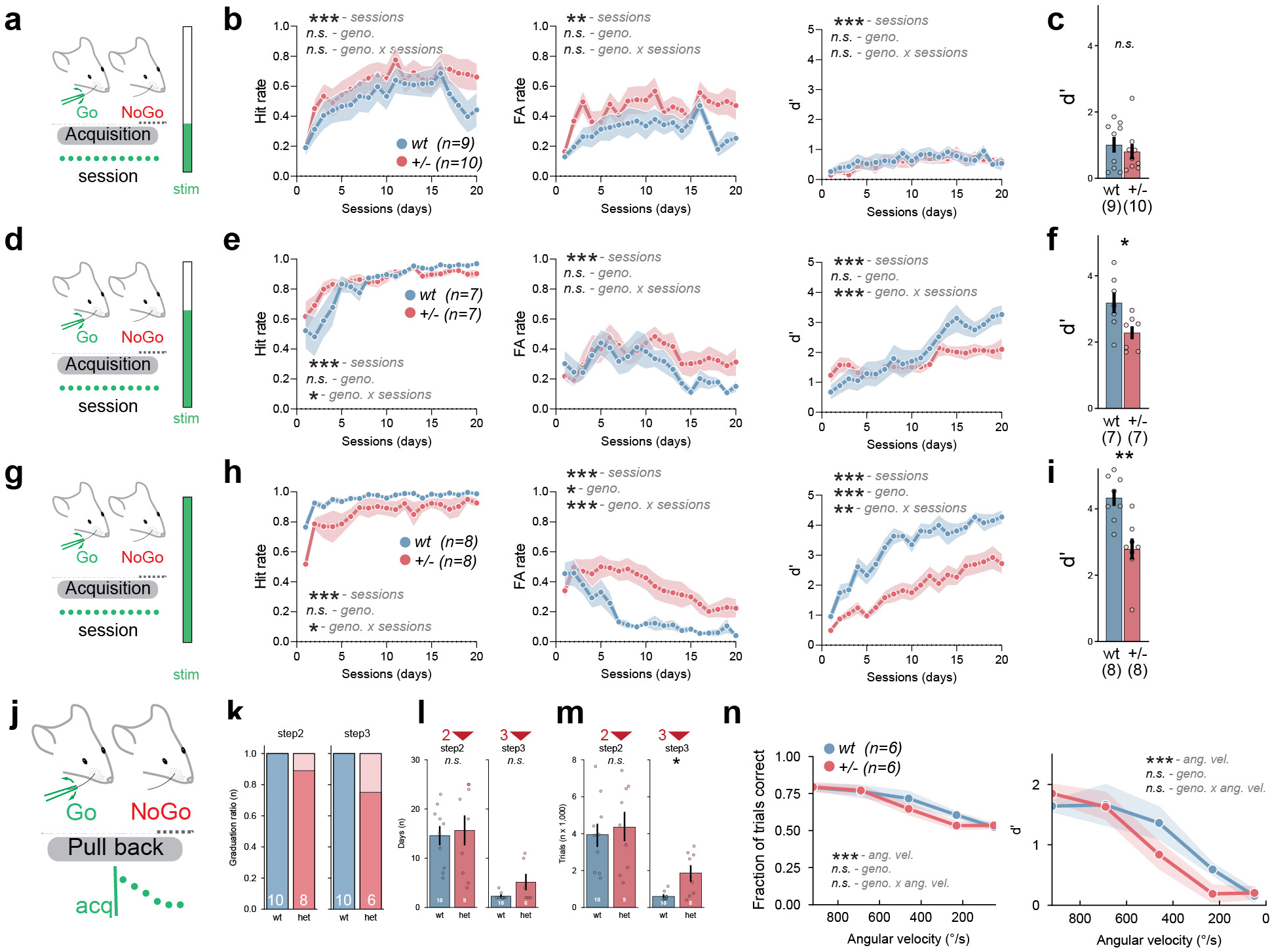
Additional measures during passive tactile stimulus (WDIL) in *Syngap1^+/−^*mice, related to Fig. 1. Behavioral measures during WDIL training for the low (400 °/s) **(a-e)**, medium (650 °/s) **(f-j)** and high (900 °/s) **(k-o)** whisker stimulation corresponding to Fig.1b. Hit/false alarm (FA) rates and d’ scores during task acquisition are indicated for the three intensity levels **(b, e, h)**. d’ scores after 20 days of acquisition for the 3 stimulus intensities **(c, f, i)**. **j-n.** Pull back experiments for the WDIL paradigm showing the ratio of mice graduating the task at the different step of training (step2 and step3) **(k)**, the number of sessions **(l)** and the total number of trials **(m)** to graduate step2 and step3 prior to the pull back experiment corresponding to Fig. 1. **n.** Fraction of total trials correct) rate and d’ score during decreasing whisker stimulation intensity of the pull back phase of the WDIL. Shaded areas and error bars represent the standard error of the mean (S.E.M.). p-value for main effects and interaction are indicated as n.s.: p>0.05, *: p<0.05, **: p<0.01, ***: p<0.001 detailed statistics are provided in Supplementary Table1.

**Supplementary Fig. 2 related to Fig. 3.**
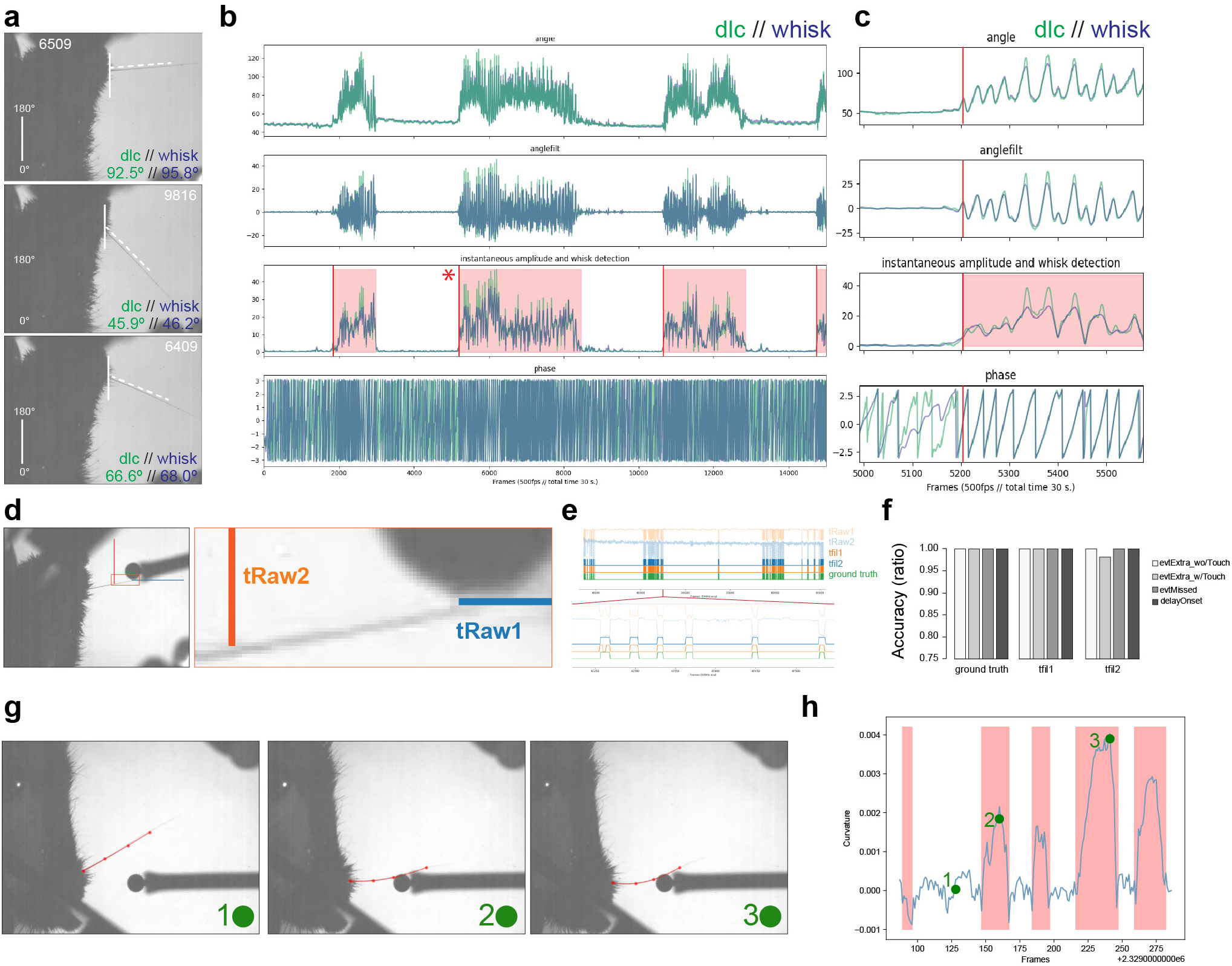
Whisker dynamics and touch detection analysis comparison between WHISK and DLC. **a.** Screen shot of 3 random frames (frame number in white) from highspeed recordings. Raw angle obtained with DLC tracking of the whisker (green) and tracking with the WHISK software (blue). **b-c.** Output of DLC and WHISK for 4 different measurements (angle, filtered angle, instantaneous amplitude and phase) during a 30 seconds period **(b)** and a snippet of data at the red star for a duration of 1 second **(c)**. **d.** Example and validation of touch detection based on thresholding and DF/F of intensity when the whisker is within the tRaw1 and tRaw2 areas. **e-f.** Comparison of filtered signal with ground truth manually analyzed data **(e)** and their relative accuracy **(f)**. **g.** Example of curvature analysis with DLC fitting a polynomial onto the first 4 proximal whisker labeled points. Green dots are references to the frame equivalent to the curvature displayed in **h**. **h.** Curvature of the whisker over time with red boxes denoting the moment of touch detected as described in **d**.

**Supplementary Fig. 3 related to Fig. 3.**
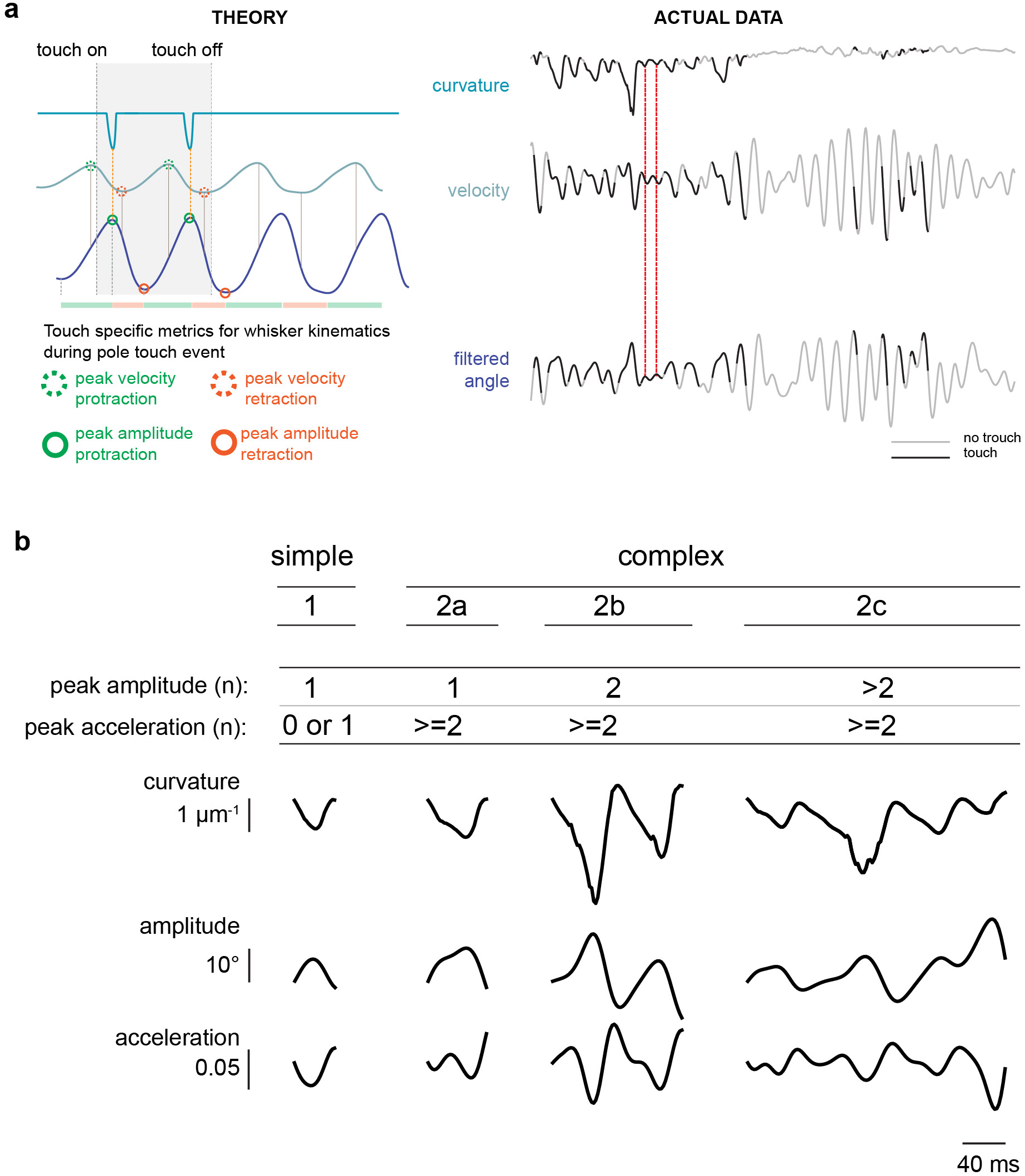
**a**. Illustration theoretical and actual data whisker kinematics during whisking and touch against a pole in a head fixed freely whisking behavior. **b.** Illustration of the categorization of different types of touches, from simple to complex with the category 2c being the most complex observed.

**Supplementary Fig. 4 related to Fig. 4.**
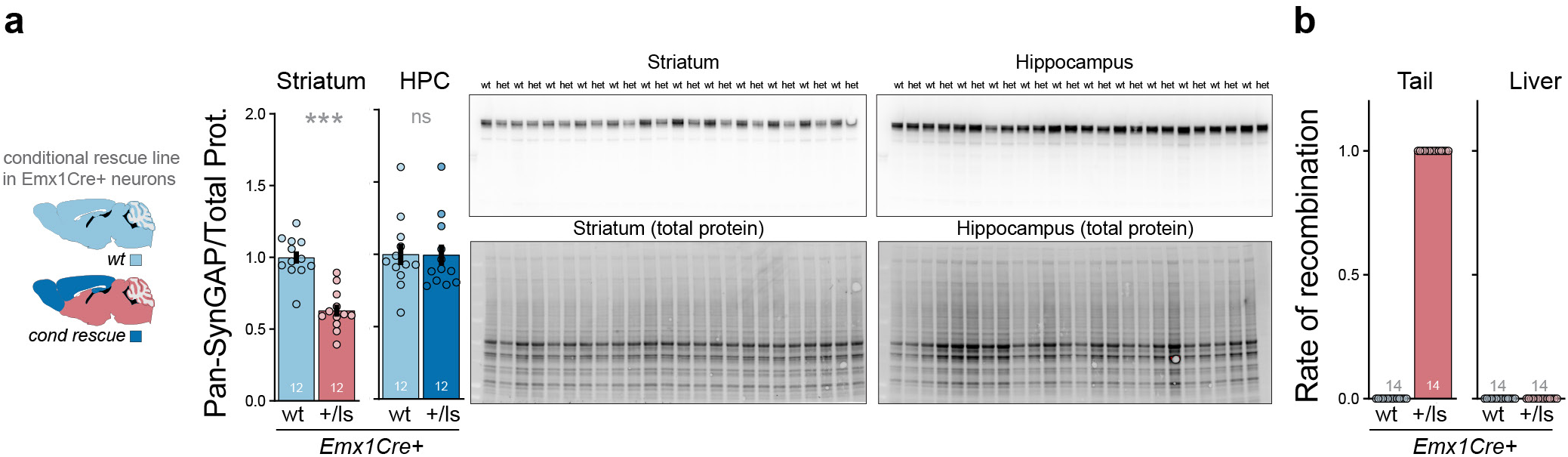
**a.** Immunoblots and quantification of Pan-SynGAP and total protein in the striatum and hippocampus from *EMX1-Syngap1-ON* mouse line. The genotype for the mouse line are reflected as such +/+ Syngap1+/+:Emx1Cre+ and +/ls for Syngap1+/lx-stp:Emx1Cre+ **b.** Rates of recombination in tail and liver tissue from *EMX1-Syngap1-ON* mouse line. The umber of animals for wild-type (wt, n=12) and *EMX1-Syngap1-ON* (+/ls, n=12) is also indicated in white within the bar graph and/or in parentheses Error bars represent the standard error of the mean (S.E.M.). p-value for main effects and interaction are indicated as n.s.: p>0.05, *: p<0.05, **: p<0.01, ***: p<0.001. Detailed statistics are provided in Supplementary Table4.

**Supplementary Fig. 5.**
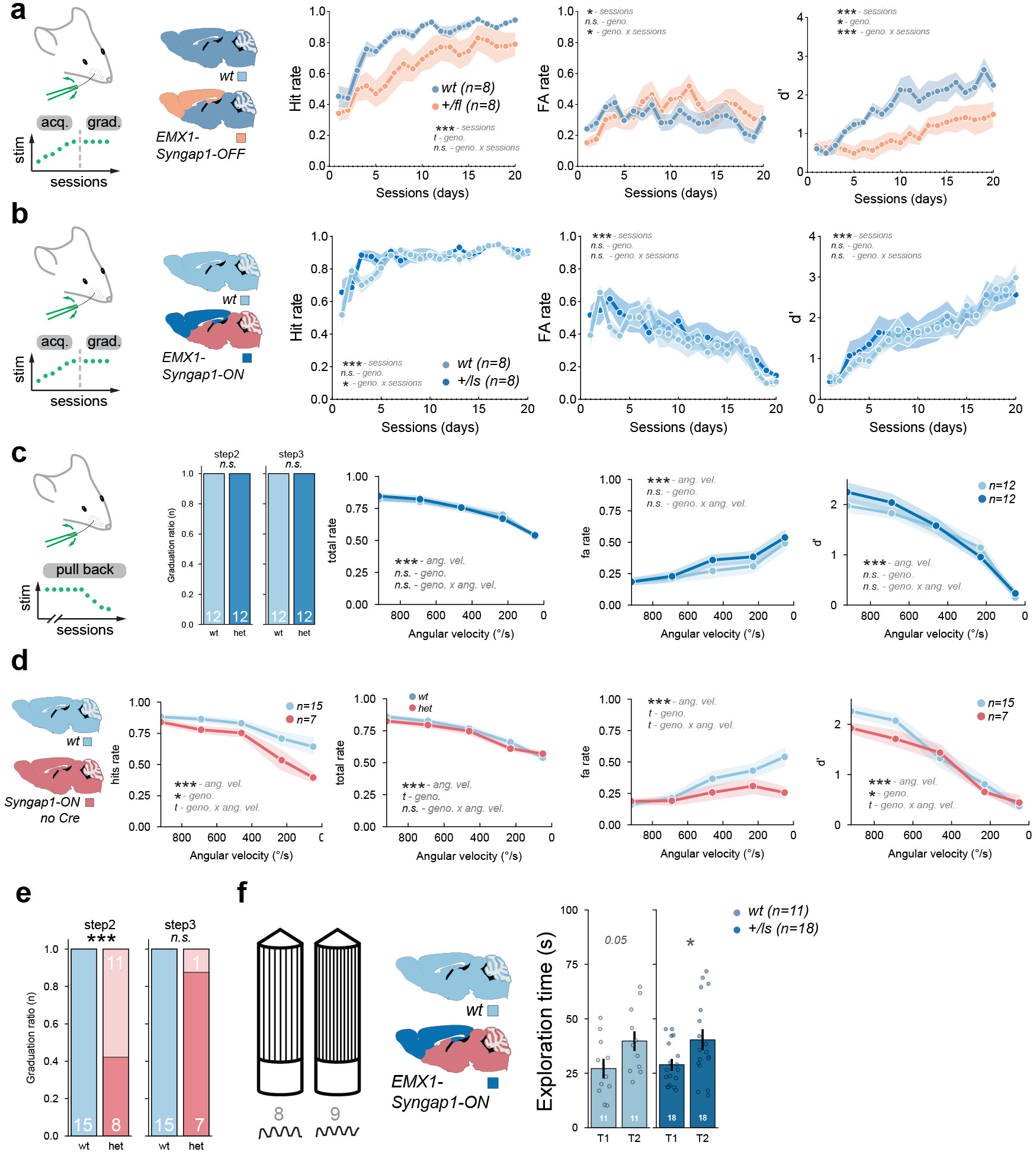
Additional measures for the modulation of *Syngap1* expression within cortical excitatory neurons during tactile sensitivity experiments related to Fig. 4. **a.** Hit/FA rates and d’ during WDIL acquisition for wt and *EMX1-Syngap1-OFF* mouse line (complementary to Fig. 4k). **b.** Hit/FA rates and d’ during WDIL acquisition for wt and *EMX1-Syngap1-ON* mouse line (complementary to Fig. 3L). **c-d**. Total fraction trials correct, FA rates and d’ in *EMX1-Syngap1-ON* mouse line (complementary to Fig. 3M) **(c)** and their genetic control counterpart which is not expressing Cre **(d)** during decreasing whisker stimulation intensity of the WDIL paradigm. **e.** Ratio of mice graduating the task at the different step of training (step2 and step3) for the genetic control counterpart of the *EMX1- Syngap1-ON* which is not expressing Cre. **f.** Exploration times during the testing phase of the NOR-T for objects with 10 groves spacing difference in *EMX1-Syngap1-ON* line (complementary to Fig. 3N). Shaded areas and error bars represent the standard error of the mean (S.E.M.). p-value for main effects and interaction are indicated as n.s.: p>0.05, *: p<0.05, **: p<0.01, ***: p<0.001 detailed statistics are provided in Supplementary Table4.

**Supplementary Fig. 6.**
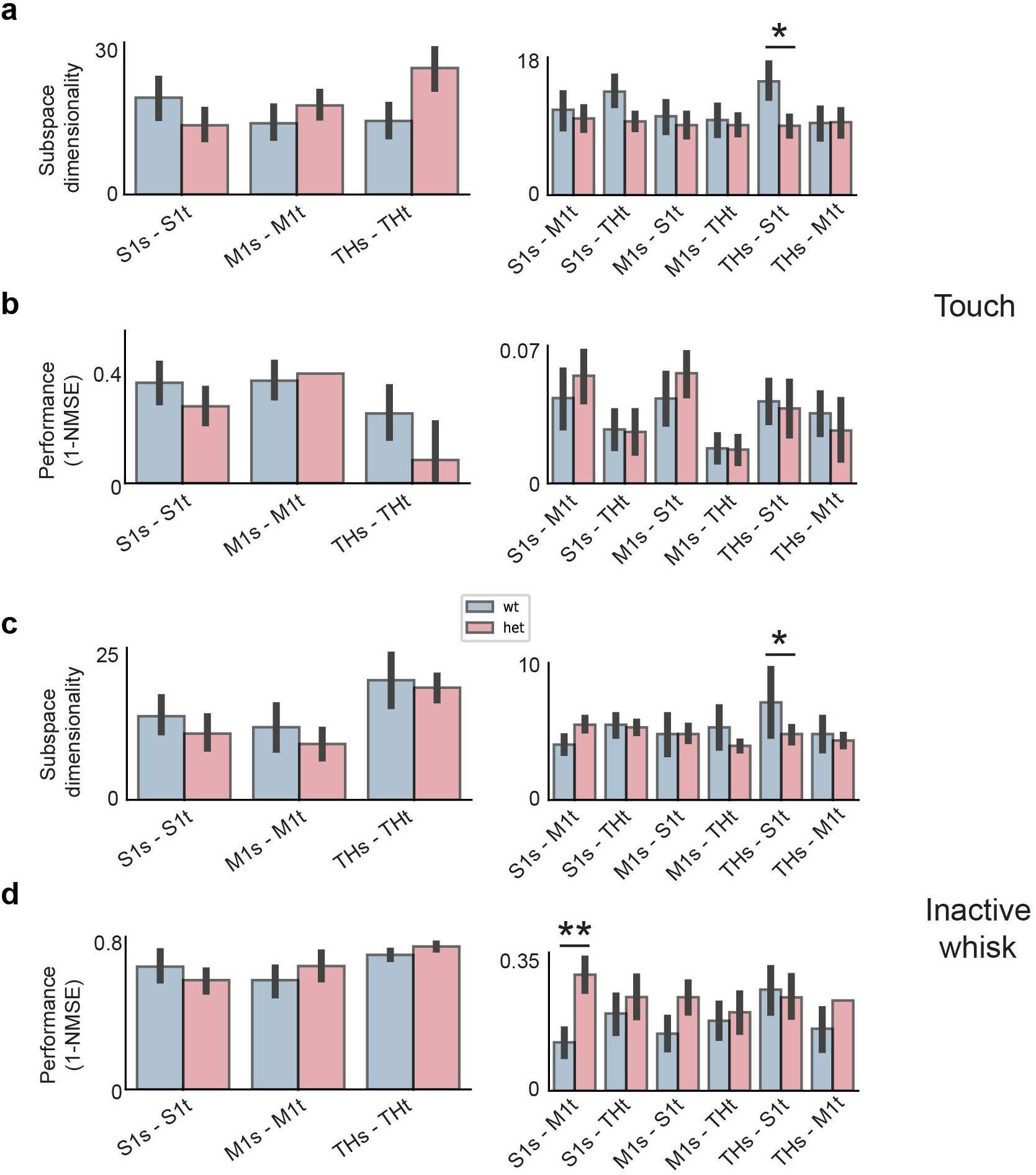
Subspace analysis of neural communication in Syngap1+/− mice across and within cortical regions during touch and inactive epochs. **a.** Subspace dimensionality for predicting neural activity in a target population from a source population, measured during touch epochs. Error bars represent SEM across different randomized choices of the source and target neural populations (see Methods). A significant increase in subspace dimensionality was observed in the TH-to-S1 pair in Syngap1^+/−^ mice compared to wild-type controls (p < 0.05). **b.** Predictive performance of target population activity based on source population activity during touch epochs. A significant increase in 1-NMSE was observed in S1-to-M1 interactions in Syngap1^+/−^ mice, suggesting that the neural activity in S1 is more predictive of M1 activity compared to wt. These results suggest altered neural communication dynamics in the S1-to-M1 and TH-to-S1 pathways. **c.** Subspace dimensionality measured during epochs during which no touches or whisks were observed. Similar to the touch epochs, subspace dimensionality was significantly increased in the TH-to-S1 projection in Syngap1^+/−^ mice compared to wild-type controls (p < 0.05), pointing to a persistent alteration in communication dynamics between these brain regions. **d.** Predictive performance during inactive epochs. A significant increase in predictive performance was observed in the S1-to-M1 projection in Syngap1+/− mice (p < 0.01), further supporting the possibility of an altered functional connectivity between these regions in Syngap1 mutants.

**Supplementary Fig. 7.**
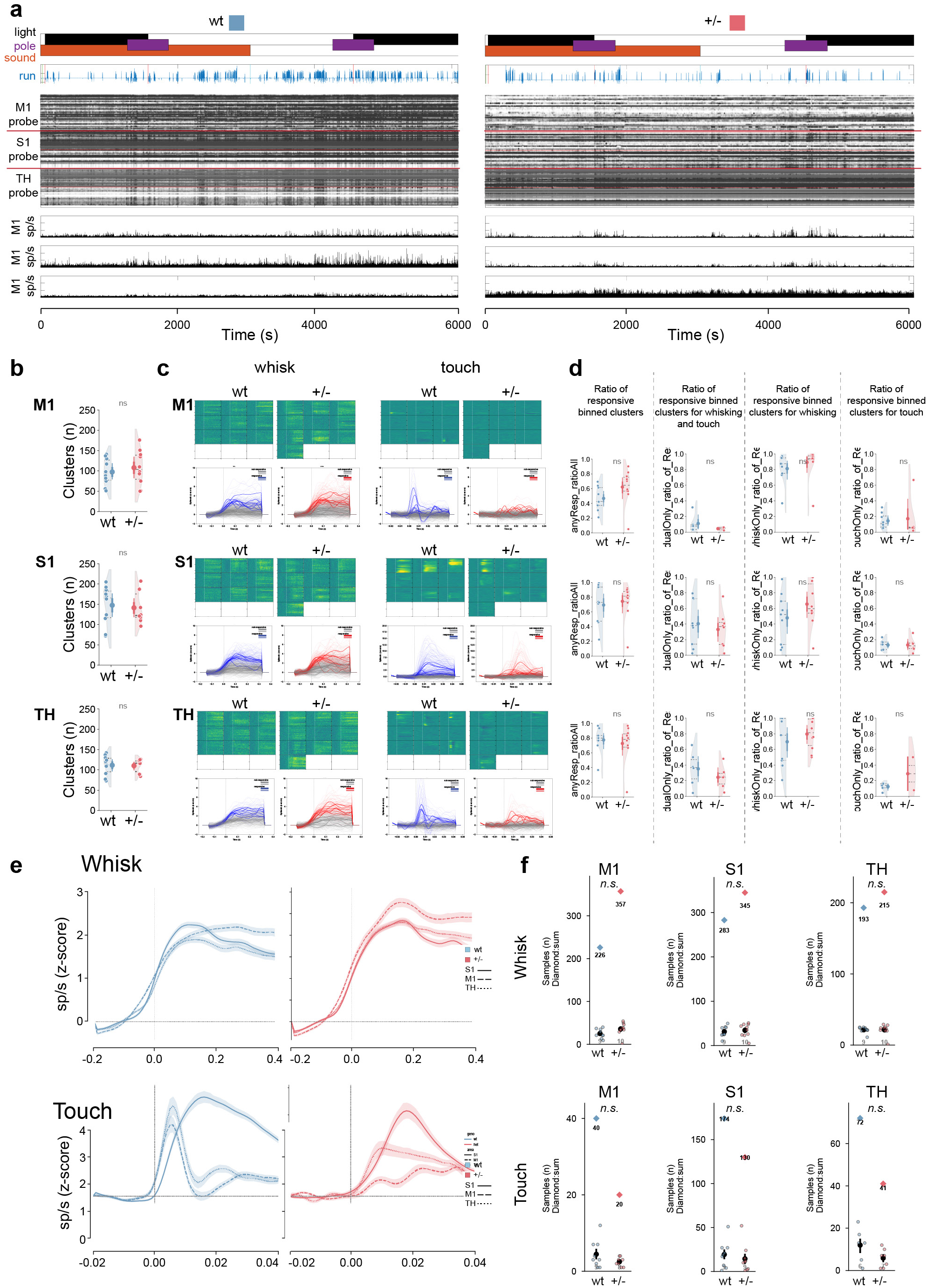
Multisite silicone probe recordings during behavior related to Fig. 6. **a.** Representation of the environmental conditions present during the recordings (dark: black, light: white, pole present: purple, sound present: orange. Motion (blue trace) on the treadmill is displayed along with Drift map and spiking activities across M1, S1 and TH for 2 representative animals (wt and het). **b.** Number of clusters detected during the recording along the three areas per animals. **c.** Raster of the PSTH analysis by depth of the neuronal activity for M1, S1 and TH. Traces for responsive (colored) and non-responsive (grey) MUA clustered by depth are displayed for wt (blue) and het (red). **d.** Quantification of the ratio of responsive clusters and their proportion relative to being responsive only to touch, whisking or both whisking and touch. **e.** PSTH of whisking and touch responses across M1, S1, and TH. **f.** Number of samples represented per animals (circles) and the sum of all the responsive traces (diamond) for wt (blue) and het (red) mice. (wt (blue) and *Syngap1^+/−^* (het, red); (n.s.: p>0.05, *: p<0.05, **: p<0.01, ***: p<0.001).

**Supplementary Fig. 8 related to Fig. 6.**
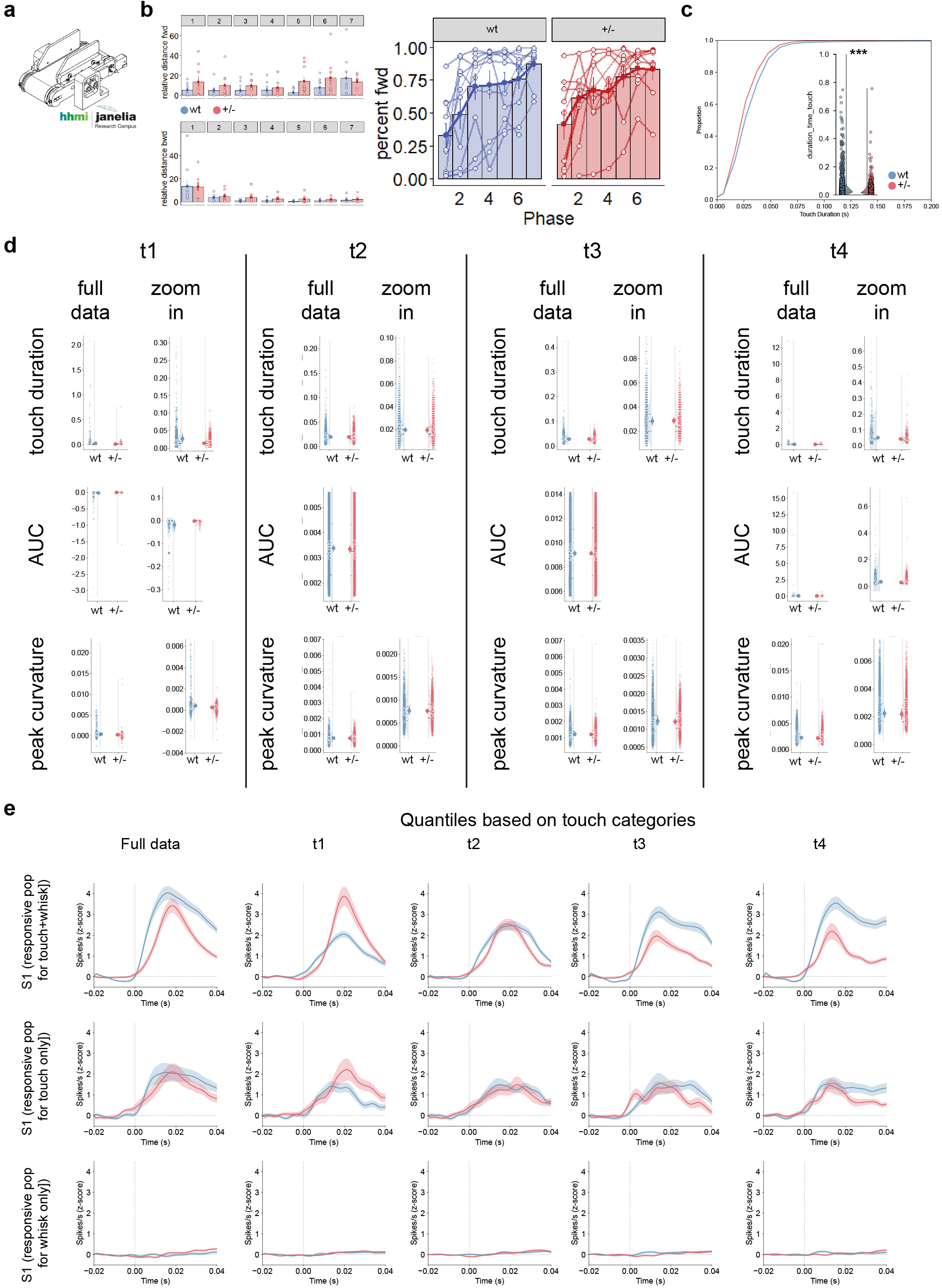
**a-d.** Behavioral metric during electrophysiological recordings. Illustration of the Janelia, Low Friction Rodent-Driven Belt Treadmill used during electrophysiology recordings **(a)**. Relative distance moved forward (top) and backward (bottom) during habituation for the first 6 days and recording (last day), the percent of forward movement during each phase of the experiment for wt and het mice **(b).** Cumulative distribution of touches in wt and het mice **(c). d.** Behavioral touch data which have been subdivided into 4 equal bins based on quartile of curvature and the corresponding measures: touch duration, AUC or peak curvature (rows) for each quartile are displayed. **e.** Classification of responsive population for whisking and touch behavior and their spike rates when subdivided by the curvature quartile described in d.

**Supplementary Fig. 9 related to Fig. 7.**
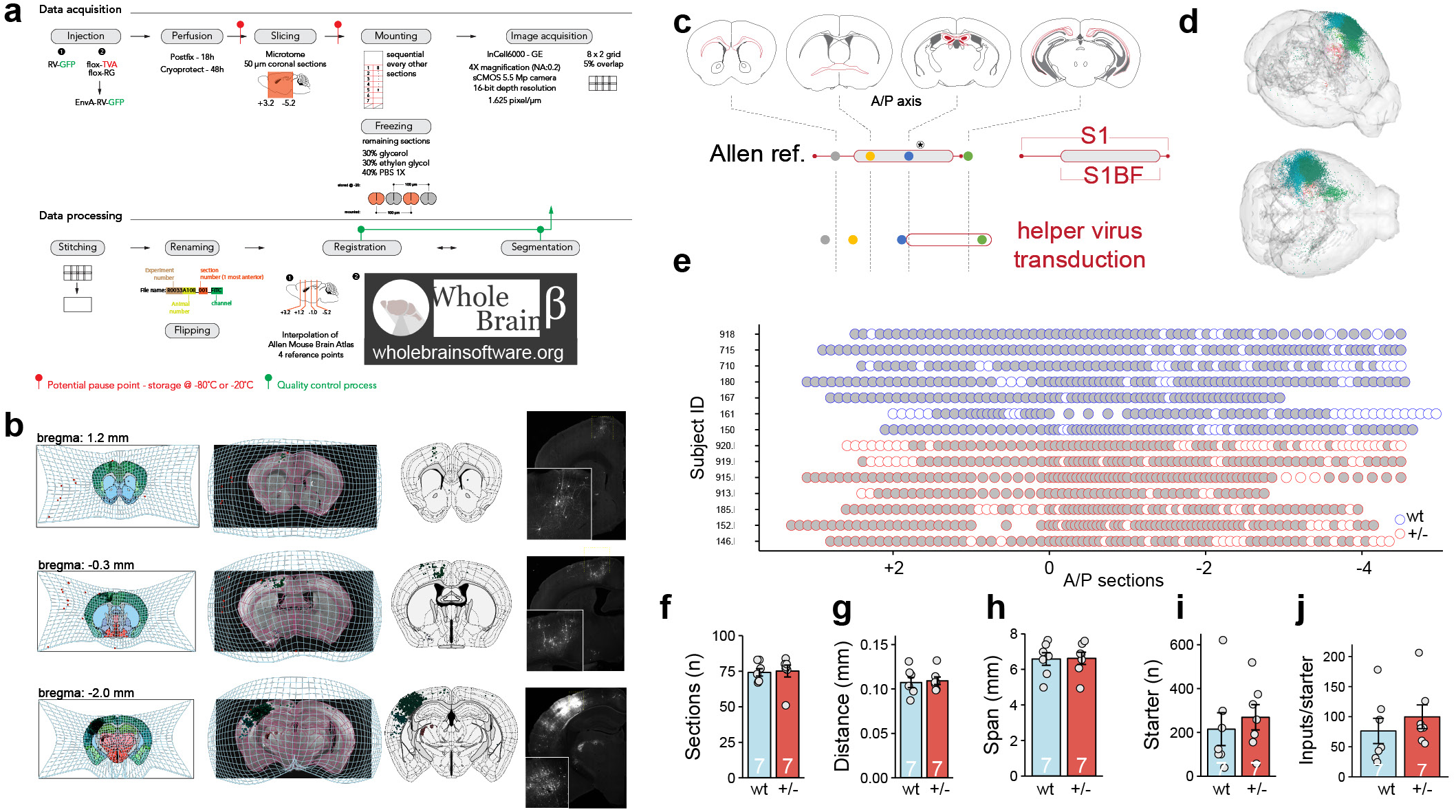
Experimental protocols related to rabies virus tracing and data associated with Fig. 5. **a.** Experimental setup and timeline for data acquisition and processing of whole brain tracing data **b.** Cellular segmentation and registration using Wholebrain software for 3 representative coronal sections. **c.** Schematic representation of viral transduction. **d.** Example 3D brain of cellular registration from 2D coronal sections. **e.** Mapping of the distribution of the coronal section used for the analysis of all the samples in the antero-posterior stereotactic coordinates. **f.** Total number of sections in which cellular count and brain region registration was performed. **G-h**. Distance between every section sampled in the study **(g)** and their brain coverage **(h)**. **i.** Number of starter cells, cells expressing both GFP and mCherry in subsamples section. **j.** Ratio of the total number of inputs to the number of starter cells counted. Error bars represent the standard error of the mean (S.E.M.). p-value for main effects and interaction are indicated as n.s.: p>0.05, *: p<0.05, **: p<0.01, ***: p<0.001 detailed statistics are provided in Supplementary Table 5.

## METHODS

### Mice

All mouse procedures were conducted in accordance with the NIH Guide for the Care and Use of Laboratory Animals, and all methods were authorized by the Scripps/UF Scripps Biomedical Research Institutional Animal Care and Use Committee. Both males and females (M/F) were used in all experiments except when explicitly noted. The design and maintenance of constitutive and conditional *Syngap1* lines have been described previously ^66,90^. Briefly, we used inbred constitutive heterozygous *Syngap1* knock-out mice (*Syngap1^+/−^*), conditional knock-out (*Syngap1^+/fl,^* JAX: #029303) and conditional rescue (*Syngap1^lx-st^* JAX: #029304) mouse lines. Emx1-Cre (JAX: #005628) mice were purchased form Jackson and crossed with *Syngap1^+/fl^*for conditional knock-out or *Syngap1^lx-st^* for conditional rescue experiments. Rbp4Cre mouse line (MMRRC_031125-UCD) was obtained from MMRC and was crossed to *Syngap1^+/−^* to study structural connectivity (monosynaptic tracing). Thy1-ChR2-YFP mouse line (JAX: #007612) was crossed to *Syngap1^+/−^* for functional validation and electrophysiological recordings. Cohort construction was designed to generate comparable sample sizes between genotypes and sexes, by allocating equal (if feasible) number of age-matched littermates from separate litters, usually more than two. Then, animals were assigned a number to hide the identity of genotype and/or group assignment. Experimentalists were blind to genotype at the time of data acquisition and analysis. Data collection occurred from mice >8 weeks of age. Mice were housed 4-5 per cage on a 12-hour normal light-dark cycle. In experiments requiring head-fixation, mice were transferred to a reverse light-dark cycle 2-3 weeks prior to headpost surgeries. Following headpost surgeries, animals were singly housed, with the addition of environmental enrichment in the form of a plastic running wheel (Bio-Serv) or a cardboard hut. Only animals that died, became non-responsive or did not participate in behavioral tasks during the study or data collection procedures were excluded from analysis.

### Headpost surgery

Headpost surgeries were completed according to established procedures with minor modifications ^23^. A custom titanium headpost was implanted onto the skull of 8-10 week old mice. Animals were anesthetized with isoflurane (5% induction, 1.5-2% maintenance) via a low-flow vaporizer (Somno Low-Flow Vaporizor, Kent Scientific) and placed into a stereotaxic frame (David Kopf Instruments). Body temperature was maintained at 37°C by a regulated pad with temperature feedback under the animal and ophthalmic ointment (Artificial Tears, Akorn) was placed onto the eyes for lubrication. The scalp was sterilized with alternating swabs of Betadine and 70 % ethanol. A small flap of skin was removed over the midline exposing both Lamda and Bregma, and the periosteum was gently cleared with a cotton swab. The skull was scraped with a scalpel and a thin layer of glue (Vetbond, 3M) was applied to the surface, reaching the wound margins. The headpost was lowered and affixed onto the skull via dental cement (Metabond, Parkell). Animals were injected (SubQ) with a cocktail of carprofen (10 mg/kg, Zoetis) and enrofloxacin (5 mg/kg, Norbrook), made in sterile saline (0.9 % NaCl, Vetivex). Animals recovered on a heating pad before being placed into their home cage. The same drug cocktail was injected once daily for the following two days for pain management and were routinely monitored for distress.

### WDIL paradigm

#### Apparatus

The whisker dependent-instrumental learning (WDIL) paradigm was performed as previously described with minor modifications ^23^. Briefly, mice were singly housed in a reverse light-dark room following headpost surgery and placed on water restriction (1 mL/d; food ad libitum), 3-7 days following recovery from headpost surgery. Animals were trained one session/day (∼5d/week), all during the dark phase of the light cycle. The behavioral rig was controlled by BControl software (C. Brody, Princeton University) running in Matlab (2013B, Mathworks) on a master PC (Dell) and a Real-Time Linux State Machine (RTLSM). The behavioral rig consisted of a light and sound proofed box constructed from aluminum rails for the frame and black hardboard (Thorlabs) with sound attenuating foam. Head-fixation parts were custom built (Max Planck machine shop) and purchased from Thorlabs.

#### Habituation

Habituation to head-fixation commenced with handling of mice for 1 d, then presentation of a custom-built stainless steel body tube for 1 d. Mice were then exposed to head-fixation for 3 consecutive days with increasing time spent under head-fixation (10, 30 and 45 mins). Mice were continuously monitored via IR light and videography (Raspberry Pi HQ; Model 3B). White noise (70 dB) was continuously played within the apparatus and all subsequent sessions to attenuate room noise. Following habituation to head-fixation, all but one whisker on each side, C2, were trimmed under light (2%, isoflurane) anesthesia and kept trimmed throughout the experiment. The following day, mice learned to associate water availability by licking water from a lickport. Detection of licks was performed electronically^91^ and precise water delivery (8 µL/ reward) was controlled with a solenoid valve and controller (INKA2424212H VHS-24V and IECX0501350A, The Lee Company). Lickport training lasted for a maximum of 10 mins/session or the total consumption of 1 mL of water per session (whichever came first), for two sessions. A lick would induce water delivery which could be consumed, however another lick would not deliver more water until a 1.5 s epoch passed. During lickport training, the C2 whisker was inserted into a plastic tube attached to a piezo to habituate the animals for training.

#### Training

Mice proceeded to WDIL training which was designed as a “detection” task, whereby the C2 whisker on the right side was plugged into the piezo actuator, acting as the Go signal and a “dummy” piezo was placed beside the whisker deflecting piezo but was not attached to a whisker, acting as the NoGo signal. Trials consisted of 50% “Go” and 50% catch (NoGo) trials in a random fashion, with the exception of no more than three consecutive trials could be of the same type. For Go trials, the whisker was deflected by the piezo actuator controlled by a linear voltage amplifier (E-650.00 and PL140.11, Physik Instrumente or EPA-008-1 – 1 and Q220-A4-203YB - 5+, Piezo.com) and a waveform generator (4054B, BK Precision), for 0.5 or 1.5 s (depending on the stimulus intensity) with a 40 Hz sinusoidal wave (rostral to caudal, 2-6 ° depending on the stimulus intensity). Bending of the piezo was calibrated using a laser-based displacement device (LD1610-0.5, Micro-Epsilon). For NoGo trials, the dummy piezo was stimulated in the same fashion as the whisker deflecting piezo, however no whisker stimulation was provided. The response window opened 0.1 s following stimulus onset and lasted for 2 s. During the response a window, a lick on the lickport resulted in a “hit” on Go trials and triggered an 8 µL water reward, and a “FA” on NoGo trials. Withholding a lick on Go trial resulted in a “Miss”, while no licking on NoGo trials resulted in a “correct rejection”. No water reward was provided on correct rejection trials and no punishments were given on Miss or FA trials. The intertrial interval remained constant at 4 s, but mice were required to withhold licking for 1.5 s before the piezo was stimulated for trials to proceed and therefore, provided some level of randomness of trial timing. Mice performed the task until satiated. Animals were trained for 20 consecutive sessions. Performance was based on a number of factors including total trials correct, discrimination index (d’; calculated as d′ = z(hit) – z(FA), with z scores computed using the function NORMSINV in excel) and Hit and FA rates. Mice were scored to be expert learners (ie. reached learning criteria) when the following metrics were achieved for two consecutive days: d’ ≥ 1.1, total trials correct ≥ 70%, Hit rate ≥ 70% and FA rate ≤ 30%. Mice that reached criteria were graduated to a reduced stimulation protocol after 20 sessions that consisted of a similar task structure however, the stimulus intensity was reduced on consecutive days (6 °, 4.5 °, 3°, 1.5°, 0.5° for angular deflection).

### NOR-T

Novel object recognition (NOR) and novel object texture discrimination (NOR-T) paradigms were developed for use with high-speed videography and conducted with *Syngap1^lx-st^* (heterozygous KO) and *Syngap1^lx-st^*x Emx1-Cre (conditional rescue) mouse lines to assess recognition memory and whisker-dependent texture discrimination in a freely moving/non-head fixed behavioral setting as a proxy to assess somatosensory cortical function in these mice.

#### Apparatus

The apparatus was assembled using infrared-transmissible plexiglass sheets (Part # ACRY31430, ePlastics) to fashion an open-top box (44(L) x 44(W) x12(H) cm). Four infrared lamps were attached at each corner, a monochrome camera (A1300, Basler) with a 25mm lens positioned 56 cm over the center floor of the box and two high-speed cameras (Spark SP-5000M-CXP2, JAI) suspended 22 cm over each of the two objects were used for video recording. The apparatus was illuminated with diffuse LED lights from above at ∼ 85 Lux to promote exploration. The Basler camera was set to record at 30 fps to assess full arena activity. The two high speed cameras were set to record at 160 fps to determine when and how long the mouse explored either object with its whiskers. The Basler camera video data was fed into Bonsai (https://bonsai-rx.org/) to track animals in real time and produce triggers in response to animals entering ROIs surrounding the objects. These triggers were fed, via an Arduino (Uno R3, Arduino), into the high-speed camera trigger to acquire frame captures. Video recordings from the high-speed cameras were recorded to a DVR system (DVR Express Core 2, IO Industries) using Coreview software for offline analysis.

#### NOR-T

3D-printed white “cog-wheel” columns (7.4 (H) x 3 (D) cm) with different numbers of teeth (50, 75 or 85 per objects corresponding respectively to 5, 8 and 9 ribs/cm) with smoothed cone tops and a separate smooth-surfaced circular base (1.8 (H) x 3 (D) cm) into which the poles screwed served as objects for texture discrimination. Bases (equidistant from the corners and 9cm from each side of the arena) were fixed throughout the task while familiar (8 ribs/cm) and novel (5 or 9 ribs/cm) poles could be interchanged throughout the task for an entire cohort. Mice were run initially in two 10 min habituation sessions with no objects in the arena. The first session conducted with only the Basler camera suspended above the arena, and the second session conducted with all three cameras positioned as in the training and testing phases. On the following days, each mouse was run in a training session with two identical familiar poles and a testing session (15 mins each) with one of the familiar poles and a novel pole separated by a 5 min intertrial interval in the home cage. Poles were cleaned with 70% EtOH, dried, and stored in clean bedding between sessions while affixed bases were wiped cleaned and dried during ITIs with urine and fecal boli removed from the arena without additional cleaning. Arenas and poles were cleaned between animals. Positions of the novel pole were counterbalanced throughout the cohort. Overall activity was assessed during habituation phases. Time spent with familiar and novel poles were compiled using session videos analyzed manually with BORIS software (https://www.boris.unito.it/) by scoring the time when animal whiskers were in contact with the objects. Comparisons between training and testing phases were performed using two way ANOVA analyses, and % novel exploration ((novel time/( novel and familiar times) x 100) and discrimination index values ((novel-familiar times/(novel+familiar times) were calculated from the pole duration data for each mouse with genotype differences assessed with unpaired t-tests. Mice with at least 10 sec of cumulative pole exploration were included in statistical analyses. For the NOR-T experiment with 8 and 9 ribs/cm object with whisker and no whisker in wt mice. For wildtype mice without whiskers 10 mice were included in the analysis and meeting the criteria out of 11 mice. For wildtype mice with whiskers 10 mice were included in the analysis and meeting the criteria out of 10 mice. For the NOR-T experiment with 8 and 9 ribs/cm object Syngap1+/−. For wildtype mice 8 were included in the analysis and meeting the criteria out of 10 mice. For Syngap1+/− mice 6 were included in the analysis and meeting the criteria out of 12 mice. For the NOR-T experiment with 5 and 8 ribs/cm object Syngap1+/−. For wildtype and Syngap1+/− mice 10 were included in the analysis and meeting the criteria out of 10 mice, respectively.

#### NOR

Novel object recognition sessions commenced once all mice of a particular cohort finished NOR-T sessions and were conducted in the same manner as the NOR-T sessions with no initial habituation sessions. Two different types of objects (two identical master locks for the familiar objects and a mini stapler for the novel object) were used in this paradigm. These objects have been verified extensively as approachable with no significant biases for exploration time in the Frick lab ^92^ and in unpublished data from the lab. The objects were temporarily fixed to the floor of the arena with heavy duty double-sided tape not accessible to the mouse. Objects and arena were managed within and between training and testing phases as in the NOR-T paradigm including counterbalancing of the novel object position. Data were subjected to the same analyses as in the NOR-T paradigm. Mice with at least 30 sec of cumulative object exploration in the NOR were included in statistical analyses. For the NOR experiment 12 wildtype mice were run in the study and 7 out of 12 met the exploration criteria, 13 *Syngap1^+/−^* mice were run in the study and 10 out of 13 met the exploration criteria.

### Free-whisking and active touch paradigms

Following headpost surgery, habituation to head-fixation and whisker trimming (described above), whisker movements (C2 whisker on the left side) were recorded during “free-air” trials (no presented) in a dark, sound-isolated chamber while head-fixed. Videos (50 s in duration) were recorded at 500 Hz from above at 640 x 480 pixels resolution with a high-speed camera (DR1-D1312-200-G2, Photon Focus) coupled with a 0.243X bi-telecentric lens (MVTC23024, Thorlabs) and Streampix software (Version 8, Norpix). The field of view was illuminated from below with an array of infrared light-emitting diodes (B001BC52W2, Amazon) with a diffusion sheet (3026, Rosco) placed above the array.

For active touch experiments, a vertical metal pole (2 mm in diameter), was moved into the whisking range of the C2 whisker (left side) via a set of feedback controlled linear actuators (L16-P 50mm, Actuonix), controlled via an Arduino (Uno R3, Arduino) and a motor control board (Part # 1438, Adafruit). Placement of the pole was adjusted manually for each animal such that the pole resided in line with the snout (rostrally) and 5-8 mm lateral of the whisker pad. The pole was presented to the animals for a total of 5 mins each, however only the first 30 s following the first touch was used for further analysis. Analysis of videos comprised of tracking whiskers offline using the Janelia Whisker Tracker ^93^, which supplied whisker traces in 2-dimensional space, followed by manual curation. Processing of this data was completed in MATLAB (2018b, Mathworks) using established protocols ^94,95^. Briefly, instantaneous phase, amplitude and setpoint were acquired by using the Hilbert transformation of the band-pass (4-30 Hz, Butterworth) filtered angle. Instantaneous frequency was obtained from the derivative of the instantaneous phase following unwrapping and conversion to whisk cycle. Angular velocity and acceleration were quantified by taking the first and second derivatives of the smoothed (Savitzky-Golay filter; 3^rd^ order, 9 frames) angle. Protraction and retraction values were resolved by obtaining positive and negative peaks of the resulting traces in question. The moment and duration of touch was determined manually via BORIS software by experimenters blinded to genotype.

### Pole localization task

Mice were trained in a pole localization task (Go/NoGo), based on previous studies (O’Connor et al., 2010), in the same apparatus and with similar pre-training methods (surgery, water restriction, habituation, lickport training) described above for the WDIL paradigm with minor modifications. This task was designed as a “discrimination” task. Briefly, mice used the C2 whisker to discriminate between two pole locations. A smooth pole (1.6 mm in diameter) was positioned (8-12 mm lateral from midline) in a home position via a high resolution and repeatable stepper linear actuator (NA11B30-T4, Zaber) coupled to a low friction linear slide (6203K317, McMaster-Carr) prior to trial initiation. Mice were required to withhold any licking for 1.5 s prior to trial initiation for the trial to proceed. On Go trials, the pole was positioned in a posterior position (3 mm from home) and lifted into the whisker range by a pneumatic linear slide (SLS-10-15-P-A, Festo) attached to the linear actuator. On NoGo trials, the pole was moved to an anterior position (3 mm from home) and lifted. An auditory tone to cue trial initiation was not used, however, the pneumatic lift generates a sound that is not covered by the white noise. Therefore, the offset of the Go and NoGo was 6 mm, but adjusted for each animal so the home position was in line with the snout. It took ∼500 ms for the pole to move into position and ∼200 ms for the pole to move upward into whisker range. During this time, mice could lick the lickport without any effect on trial outcome. The response window started ∼1 s after the start of the upward pole movement and was open for 2 s. Mice made their decision during this time by licking an electronic lickport. On Go trials, if the animal licked, they received an 8 µL water reward and was considered a hit. If they withheld their lick, it was considered a miss. On NoGo trials if the animal licked, it was considered a FA and the animal received a 15 s timeout with the pole remaining in the upward position. If they withheld their lick, it was considered a correct rejection, however no reward was provided. Following the end of the response window (and punishment time), the pole dropped and was moved back to the home position and a 4-6 s intertrial interval began. Animals performed 200-300 trials per session, 1 session/day, ∼5 session/week for 29 sessions. Trial types (Go/NoGo) were presented randomly with the only limitation of no more than three of the same trial types could be presented in a row. Animal performance was quantified in a similar fashion to the WDIL paradigm described above.

High speed videography of mouse whiskers was performed on five sessions throughout the training, including Session 1, 17 and 25. These were picked to include whisking behavior when the animals were completely naïve to the task (HS1, session1), when the animals (at a population level) were well into the learning phase (HS3, session17), and at the end of training (HS5, session 25). An addition 2 sessions were recorded based upon individual learning curves. HS2 was performed for each animal when their learning curve showed a steep acceleration (session 10-15) indicating learning. HS4 was conducted after the animal had reached criterion for expert level (as described in the WDIL paradigm) for two consecutive days.

Acquisition of whisking behavior and pole touches during the task was performed using a high-speed camera (Spark SP-5000M-CXP2, JAI; ∼500 Hz frame rate, 640 x 480 pixel resolution) under infrared illumination and recorded to a DVR system (DVR Express Core 2, IO Industries) using Coreview software. Each trial triggered a new recording that extended 3 s prior and 3.1 s after upward pole movement (using a pre-trigger buffer). This provided whisker activity prior to pole movement, sampling the pole and during decision making.

Whisker tracking and processing was performed using the Janelia Whisker Tracker and custom Matlab scripts, as described above, on a trial-by-trial basis. For whisking analysis during electrophysiological recordings whisker tracking was performed with DeepLabCut where 5 points along the single whisker track the whisker location. Whisker angle was calculated from the base of the whisker and the whisker pad to the next marker on the whisker. The moment of touch was quantified using a threshold-based method assessing the DF/F of pixel intensity within three 12x2 pixel areas 1 pixel away and tangential to the pole.

### Monosynaptic tracing

Viral preparation and monosynaptic tracing were performed as previously described ^74,76^. Monosynaptic inputs onto a genetically define cell population of Layer 5 neurons in S1 were targeted with helper virus AAV9-CAG-Flex-RG (Addgene 48333) and AAV9-CAG-Flex-TCB (Addgene 48332,^75^) in conjunction with pseudotyped rabies virus EnvA-RV-GFP in Rbp4Cre mouse line crossed with *Syngap1^+/−^*. AAV9-CAG-Flex-RG (titer: 1.10^12 IU/ml) and AAV9-CAG-Flex-TCB (titer: 1.10^12 IU/ml) were mixed as a 1:1 ratio and 400 nl were unilaterally injected in S1BF (AP: -1.3 mm, ML:+3.0 mm and DV:-0.45 mm from Bregrma) at a rate of 200 nl/min. Two weeks after the first injection 400 nl of EnvA-RV-GFP (titer: 1.10^8 IU/ml) was injected in the same location at a rate of 200 nl/min. 7 days after injection of EnvA-RV-GFP the animals were deeply anesthetized and intracardially perfused with 0.1 M PBS followed by 4% paraformaldehyde.

### Histology and image acquisition

After perfusion the brains were post fixed in 4% PFA overnight, placed in 30% sucrose for 3 days prior to being snap frozen in 2-methylbutane and stored at -80 °C. On the day of slicing the dorsal part of the brain was placed on a grid, aligned and subsequently embedded in OCT against a frozen razor blade at the posterior end of the brain, creating a plane perpendicular to the dorsal part of the brain and parallel to the microtome blade. This ensures the proper alignment of the brain in the medio/lateral and dorso/ventral axes which facilitates registration to the mouse reference atlas. The whole brain was sliced on a microtome at 50 um intervals, with every other slice mounted with DAPI (P36931, Invitrogen) onto a microscope slide. The brain was subsequently imaged on an INCell Analyzer 6000 (GE) for rapid acquisition with a Nikon 4X/0.20, Plan Apo, CFI/60 at 1.625 um pixel size. Acquired images were obtained using FITC (excitation: 488nm, emission: 525nm).

#### Quantification of monosynaptic tracing

Retrograde labeled cell bodies were segmented and registered onto the mouse reference atlas using WholeBrain software ^76^. To quantify the starter cell populations, sections with red signal from the transduction of AAV9-CAG-Flex-TCB were pre-identified and re-acquired on the InCell 6000 with a Nikon 10X/0.45, Plan Apo, CFI/60 at a resolution of 0.65 um pixel size for both FITC (excitation: 488nm, emission: 525nm) and dsRed (excitation: 561nm, emission: 605). The overlapping population of green and red cell bodies were manually quantified with ImageJ and defined as the starter cell population. All of the identified inputs for any given brain regions were normalized to the total number of starter cells identified per mouse brain.

#### Quantification of electrode tracks

Sections were imaged on an IN Cell Analyzer 6000 (GE) with a Nikon 4X/0.20, Plan Apo, CFI/60 at 1.625 um pixel size with dsRed (excitation: 561nm, emission: 605) corresponding to the signal emitted by DiI to identify the electrode and FITC (excitation: 488nm, emission: 525nm) to capture the outline of the brain with autofluorescence of the tissue. The acquired images were stitched with channels merged prior to atlas registration and electrode track visualization, which was performed with SHARP-TRACK^96^.

### Electrophysiological Recordings

Mice went through a protocol to allow acquisition of electrophysiological recordings in multiple brain sites in an awake, head-fixed setting. Details of this protocol are detailed below.

#### Day1 - Surgery

The mouse underwent headpost surgery, see prior Methods, with the following addition for electrophysiological recordings. The entire scalp was removed and the periosteum was gently cleared with a cotton swab. The skull was then leveled in the antero-posterior axis by having Bregma and Lambda in the same plane, while the medio-lateral axis was leveled by adjusting the lateral point in the same plane 2 mm from the midline on each side. Enough bone was shaved from the skull with a 0.6 mm drill bit to create four reference points. This procedure did not result in exposing the brain. The reference points were located above the future electrode point of entry to reach the motor cortex (M1; AP:1.0, ML:- 1), the thalamus (TH, AP:-1.5, ML:-1) and the somatosensory cortex (S1, AP: -2, ML: -3.5), along with the front left corner where the headpost would be located (AP: -5.5, ML: 1.5). A silver wire pre-soldered to a female gold pin was in contact with the brain above the cerebellum (AP:-5.5, ML:0). Dental cement (Metabond, Parkell) was applied to secure the 3D printed plastic well, headpost and ground wire. At the end of the procedure Kwik-Cast (World Precision Instruments) was applied within the well to protect the skull surface for downstream procedures.

#### Day4 - IOS imaging

Three days after recovery of the headpost surgery, Intrinsic Optical Imaging (IOS) was performed to measure the hemodynamic response of the somatosensory cortex to define the cortical area corresponding to brain activity responses after single whisker stimulation (C2). The mouse was anesthetized as described in the headpost surgery protocol. All the whiskers were fully trimmed to the base of the whisker pad except for the whisker C2 contralateral to the cortical area of interest (trimmed to a ∼5 mm length). The skull was drilled in concentric circles over S1 (AP: -2, ML: -3.5) through the spongy bone. Debris were removed with compressed air and Ringers solution was applied to cool the bone and remove the debris. When cerebral blood vessels became visible a scalpel blade (#501251, World Precision Instruments) was used to shave the bone further until blood vessels were clearly visible. The mouse was moved from the stereotaxic instrument to the IOS imaging rig and the anesthesia was reduced to 0.7% in conjunction with the injection of a sedative (chlorprothixene; 1 mg/kg, intramuscular). Kwik-Cast was removed from the well and ophthalmic ointment (Artificial Tears, Akorn) was applied on the edge of the well filled with saline and sealed with a glass coverslip. IOS imaging was subsequently performed as previously described^23^. Briefly, imaging was performed under a 4x objective on an upright microscope (BW51X, Olympus) and the skull was illuminated with a 630 nm LED light ring mounted to the objective. Images were acquired with a Zeiss Axiocam camera (Carl Zeiss) controlled my µManager software (Open Imaging, Inc.). The C2 whisker was deflected for each IOS trial (50-70 trials total) and resulting images were processed using the IO and VSD Signal Processor plugin in ImageJ ^97^.

#### Day5-11 – Habituation

The day after IOS imaging, the mouse was habituated to head-fixation on a treadmill (https://www.janelia.org/open-science/low-friction-rodent-driven-belt-treadmill). The mouse was handled for 5 min in the morning. In the afternoon it was handled for 5 min prior to being introduced to the treadmill and allowed to explore the treadmill for 5 min. Day 6 was the first day of gradual head fixation with the animals being head-fixed for 15 min, 30 min on day 7, 1hr on day 8 and day 9, and finally 2hr on day 10 and 11.

#### Day12 – Surgery

The mouse was placed in the stereotaxic apparatus (KOPF) to perform 3 craniotomies for future electrode insertion. The C2 whiskers was trimmed to a length of 5 mm. The skull was properly positioned (correction for antero/posterior and medio/lateral tilt). Circular craniotomies of a 1.5mm diameter were drilled automatically with the Neurostar surgery robot with a 0.2 mm drill bit (Harvey tool) above the area marked during the headpost surgery and identified with IOS for the S1 craniotomy. Once the drilling was performed, the inner part of the bone was removed. Gelfoam and Ringers solution were applied to minimize potential bleeding. Kwikcast was applied to protect the brain after the craniotomy and the mouse was placed on a heating pad for recovery before returning to its home cage for 3 to 4 hrs.

#### Day12 – Apparatus

The apparatus was a 30 inx30 inx50 in custom made noise-attenuated chamber mounted on a breadboard (MB30, Thorlabs). The recording consisted of a 20 min period in the dark followed by 40 min in the light and 20 min in the dark. In the middle of this sequence white noise that was playing (70 dB) was turned off. In addition, 2 pole presentation epochs of 10 min occurred 5 min prior to the light transition. Light, white noise, and actuators (L16-P 50mm, Actuonix) for the pole presentation were controlled via Matlab through a NI DAQ (USB-6363).The signals for the NI DAQ converged to the eCuber Server (White Matter) for synchronization. The mouse was video monitored with e3Vision Cameras (White Matter). Mouse whisking and touch behaviors were acquired at 500 Hz with a high-speed video camera (DR1-D1312-200-G2, Photon Focus) and a variable zoom lens (Computar) with StreamPix 6.0 software for the entire duration of the recording. The high speed captured video was downsampled with ffmpeg software (Version 4.0.2) before data analysis. IR light illuminated the mouse and a custom IR backlight was located underneath the mouse during video recording. Three 3-axis micromanipulators (New Scale Technologies) were mounted on an inverted 360 MPM-1 platform (New Scale Technologies) to enable probe insertion at 3 different locations. H2 probes for TH, H2 or H3 probes for S1 and H3 probes for M1(Cambridge Neurotech) were connected via Molex, Omnetix connector adaptor to the HS64 head stages (White Matter), which were connected to the e3 Server (White Matter). The fully retracted probes were positioned in the insertion probe axes above predetermined stereotaxic coordinates of the area of interest. Prior to placement on the manipulators the probes were coated with DiI (Life Technologies, #V22885).

#### Day12 – Data acquisition

The mouse was placed on the treadmill and moved to the rig for probe insertion and recording. The reference and the grounds of the probes were grounded to a common ground shared with the ground of the animals. After the mouse was placed in the recording apparatus all the probes were manually lowered to a few millimeters above the skull surface. The position of the probes was monitored with a digital microscope (Dino-Lite, Premier) and the S1 probe was refined based on the blood vessel map corresponding to the responsive area of whisker stimulation determined by IOS imaging. The probes were then lowed at coarse intervals with the manipulator to break the dura. When all the probes were implanted, the probes were further inserted automatically at a rate of 200 um/min until the pre-determined target depth was reached. The probes settled for 30 min prior to recording. Electrophysiological data were acquired at 25 kHz using the Open Ephys GUI with the e3 custom module. Binaries were acquired along with digital and analog data streams from the NI DAQ box through the E3 servers. Whisking and touch behavior were acquired at 500 Hz. At the end of the recording the probes were removed and cleaned and immersed in 1 % Tergazyme overnight.

#### Data Analysis – Electrophysiology

Raw binaries were processed in Matlab (2018). Median noise filtering was applied and channels were sorted according to the channel map, followed by common average referencing. From the binaries, action potentials (APs) and local field potentials (LFPs) were extracted using a Butterworth low pass filter between 0.5 and 100 Hz for the LFPs and a high pass filter between 300 Hz and 6kHz for the APs. The filtered APs were run through spike sorting software (Kilosort 2.0). Clusters of spikes were identified as “good” isolated units and “multi-unit activity”. Spike times for these categories were assigned to a specific depth in order to obtain a measure of overall spiking at a given location. Whisking onset was defined when the whisking amplitude increased by 2° within a 250 ms period ^69^. The epochs during which the pole was present were excluded from the whisking analysis. Spike rate was aligned to the whishing onset to obtain peri-stimulus time histograms (PSTH) based on depth of the probes. The PSTH for whisking were performed with analysis window from -0.25 to 0.5 s and a baseline from -0.25 to 0 s with bins of 10 ms. The touch onset was identified with a threshold-based method registering whisker and pole interaction. Manual validation of the method was performed on a subset of the data. The pole was position was maintained between both pole presentations and kept relatively standard between animals via following the position of a guided template overlay on the live camera view. The whisker resting state was posterior to the pole and any touch occurring when the whisker was anterior to the pole was removed from the analysis. PSTH analysis for touch was performed similarly to the whisker’s PSTH with 1 ms bins, an analysis widow from -0.025 to 0.05 s and a baseline from -0.025 to 0 s. The z-scored PSTH for whisking and touch were smoothed with a Gaussian kernel using the smooth function. Artifacts at the end and beginning of the traces were removed from the analysis. Responsive clusters by depth were defined as z-score firing rate above 1.5 in the post baseline window. The Findpeak function was used to identify the peak value of the trace crossing this threshold and onset of the response was defined as the first inflection point of the trace from the identified peak to the beginning of the trace.

### In vivo single electrode field recordings

LFP recordings were made on a custom in vivo system as described previously^23^. Briefly, mice were anesthetized with 1.8 g/kg urethane (Sigma-Aldrich), followed by implantation of a custom headplate, and a 1 mm craniotomy was made over S1. The pipette was lowered 500 µm from the brain surface in S1 (AP: 3.5, ML: 2). Recordings were performed in current-clamp mode with the following internal solution in the electrode (mM): 130 potassium gluconate, 5 KCl, 10 HEPES, 10 sodium phosphocreatine, 0.4 EGTA, 1 Na-GTP and 4 Mg-ATP (pH 7.3, 285-290 mOsm). Electrophysiological signals were amplified with Multiclamp 700B (Molecular Devices), filtered at 2 KHz, digitized (10 KHz) with an NI USB-6363 DAQ (National Instruments) and recorded using the NI acquisition system in Matlab. Optogenetic stimulation was controlled via the NI acquisition system in Matlab and relayed through the NI DAQ, a LED controller (LEDD1B, Thorlabs), fiber optic LED 470nm (M470F3, Thorlabs) and a fiber optic cannula (CFML12L02, Thorlabs) inserted in M1 (AP:1.0, ML:-1, DV:-0.5). Before each experiment the power of the laser was calibrated to obtain the following powers 1, 5, 10, 15, 20, 25,30, 35 and 40 mW/mm2 at the end of the fiber optic. Piezo stimulations were performed on a single whisker with a deflection of 200 µm at 2 mm away from the whisker pad (6° or 1200 °/s). To obtain LFPs, 30 trials were averaged.

### Neural communication subspaces

To analyze neural communication subspaces in neural populations for wild-type and *Syngap1^+/−^* mice we applied the regularized regression technique described in the noted reference^71^. We analyzed neural data collected from three types of behavioral epochs: whisks, touches, and inactive periods. For each whisking or touch event, a symmetric time window was centered on the event time. For whisking, the window spanned 500 ms before and after the event, while for touch, the window extended 50 ms before and after. The inactive epochs were established by finding intervals without whisking or touch events. Intervals of at least 1 second between events were considered as potential windows of inactivity. Each window was defined to begin at least 1 second after one event and end at least 1 second before the next, ensuring that the neural activity within the window was not contaminated by any whisks or touches.

#### Data preprocessing

Spike trains collected from WT and HET mice were transformed into spike rate arrays for the different brain regions (M1, S1, and TH). To do so, we convolved spike trains with a moving window (20 ms wide), allowing for the smoothing of the spike rates over time. The smoothed data were then downsampled to reduce the time resolution, which allowed for more efficient analysis of the neural dynamics. The downsampling process was implemented with a stride of 5 ms, reducing the size of the dataset while maintaining the overall trend in neural activity. This downsampled data was then used as input for subsequent analyses such as dimensionality reduction and regression.

To enable fair comparisons of neural communication between brain regions, neuronal populations recorded in each area were matched based on their firing rates. The goal of this matching process was to control for variability in neural activity levels and ensure that equivalent neural populations were being compared across regions.

First, the mean spike rate of each neuron was calculated across all time points and event types. Neurons with low spike rates (< 0.1 Hz) were excluded from further analysis. Once this filtering step was complete, neurons belonging to one area were grouped into 10 equal-width bins according to their spike rate. The binning was conducted separately for the source and target neurons, resulting in a bin index for each neuron in the source area *X* (b_X_(i), for i = 1, …, N_X_) and the target area *Y* ((b_Y_(j), for j = 1, …, N_Y_). Within each bin, neurons from the source and target areas were randomly paired. If the number of neurons in the source area within a particular bin N_X_ was larger than the number of neurons in the target area N_Y_, or vice versa, the smaller of the two numbers was chosen so the number of neurons in each population would be matched. This random matching procedure was repeated 25 times for each pair of populations. When source and target dimensions belonged to the same brain region, neurons within the region were randomly split into two groups to serve as source and target populations. We then followed the same matching procedure as for cross-area comparisons.

#### Reduced Rank Regression with Ridge Regularization

After the neurons were matched across regions, reduced rank regression (*RRR*) was employed to analyze communication between the source and target areas. The relationship between the activity in the source population X and in the target population Y was modeled as:

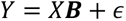

where *X ∈ R^n×p^* represents the neural activity in a source population containing *p* neurons, *Y ∈ R^n×p^* represents the neural activity in the target population with the same number of neurons, *B ∈ R^p×p^* is the matrix of regression coefficients, and ɛ is the noise term. To identify the most important communication dimensions we decomposed B into lower-dimensional components:

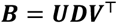

where *U ∈ R^p×r^* and *V ∈ R^p×r^* are orthogonal matrices and *D ∈ R^r×r^* is a diagonal matrix containing the singular values. The rank *r* was selected based on cross-validation as described below. To reduce overfitting, **B** was determined by applying a ridge regression model which minimized the following objective function:

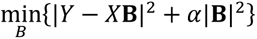

where *α* is a regularization parameter which controls the magnitude of the penalty on large coefficient values. To determine the optimal value of *α*, a grid search was conducted over 100 logarithmically spaced values from 10^-5^ to 10^5^. The best *α* was chosen as the largest *α* within one standard error of the best-performing model across all *α* values using cross validation, ensuring that the model is regularized while maintaining predictive accuracy. The rank of the reduced regression model was varied from rank 2 to full rank, which varied depending on the number of neurons obtained after the matching procedure.

#### Cross-Validation and statistical testing

To assess the performance of the RRR models at different ranks, 10-fold cross-validation was used. In each fold, the dataset was split into training and testing sets, with the RRR model trained on the training set and evaluated on the test set. We measured each model’s performance using including the Normalized Mean Squared Error (NMSE) calculated as:

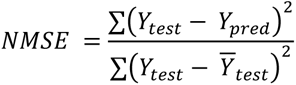

where *Y_test_* is the observed activity in the test set, *Y_pred_* is the predicted activity, and 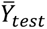 is the mean of the test set. Cross-validation was performed for each rank, generating average performance metrics and standard errors across 10 folds. The optimal rank was chosen as the smallest number of dimensions *r* where the 1-NMSE remained within one standard error of the mean from the best model. To validate the robustness of the observed genotype effects, Monte Carlo permutation tests were conducted. The genotype labels of the matched neuron pairs were randomly shuffled, and the RRR models were re-fitted to generate a null distribution of performance metrics. The observed difference between WT and HET mice was compared to this null distribution to calculate a two-sided p-value. Each permutation test was repeated 10,000 times. A *Mann-Whitney U test* was used to compare the distributions of 1-NMSE scores and SEM ranks between WT and HET mice. To account for the large number of statistical comparisons, the *Holm-Bonferroni* correction was applied to the resulting p-values.

### Statistics

Data analyses were conducted in GraphPad Prism (version 9.4.1, GraphPad Software) or custom Python scripts (version 3.9). Linear mixed models with repeated measures were used in passive and active WDIL experiments to determine differences in overall genotype performances and their learning differences within these tasks by assessing interactions between genotype and session progression during acquisition phases or angular velocity degression during “pullback” phases. ANOVAs and t tests were utilized to compare genotype differences within, between, or among different categories or stages of WDIL experiments, as well as NOR/T experiments, whisker kinematics comparisons, synaptic/circuit connectivity experiments and brain region-specific neural activity experiments. Kaplan-Meier survival curves were utilized to compare genotype differences in reaching endpoints, namely, reaching criteria for a particular stage of a WDIL task. Data are presented as mean ± SEM unless otherwise noted. D’Agostino-Pearson omnibus normality test was applied to test data normality and the appropriate parametric or non-parametric statistical test was performed accordingly. The statistical tests used and number of observations are reported explicitly in a comprehensive table **(Table S1)**. *P*-values are corrected for multiple comparisons when multiple simultaneously statistical comparisons were performed. No statistical test was used to predetermine sample sizes, however, our sample sizes are similar to those previously reported in the field ^23,69^.

